# Θ-Net: Achieving Enhanced Phase-Modulated Optical Nanoscopy *in silico* through a computational *‘string of beads’* architecture

**DOI:** 10.1101/2023.01.24.525271

**Authors:** Shiraz S/O Kaderuppan, Eugene Wai Leong Wong, Anurag Sharma, Wai Lok Woo

## Abstract

We present herein a *triplet* string of concatenated O-Net (‘*bead*’) architectures (formulated as discussed in our previous study) which we term ‘Θ-Net’ as a means of improving the viability of generated super-resolved (SR) images *in silico*. In the present study, we assess the quality of the afore-mentioned SR images with that obtained via other popular frameworks (such as ANNA-PALM, BSRGAN and 3D RCAN). Models developed from our proposed framework result in images which more closely approach the gold standard of the SEM-verified test sample as a means of resolution enhancement for optical microscopical imaging, unlike previous DNNs. In addition, *cross-domain (transfer) learning* was also utilized to enhance the capabilities of models trained on DIC datasets, where phasic variations are not as prominently manifested as amplitude/intensity differences in the individual pixels [unlike phase contrast microscopy (PCM)]. The present study thus demonstrates the viability of our current multi-paradigm architecture in attaining ultra-resolved images under poor signal-to-noise ratios, while eliminating the need for *a priori* PSF & OTF information. Due to the wide-scale use of optical microscopy for inspection & quality analysis in various industry sectors, the findings of this study would be anticipated to exhibit a far-ranging impact on several engineering fronts.

## Introduction

Over the past few decades, the field of artificial intelligence (AI) has witnessed considerable progress & deployments in numerous applications, such as computer vision [1] & [2], speech & text recognition [3] & [4], as well as cyber security [5] & [6], amongst others. Fueled primarily by developments in computing hardware resources (such as GPUs & memory [7]), a particular subset of AI algorithms (termed *deep neural networks*, or DNNs in short) has played a fundamental role in driving this revolution. In this respect, it would be prudent to evaluate the impact of these DNNs in propelling developments in healthcare & ecological studies, two aspects which play prominent roles in circumventing present-day global dilemmas, such as pandemics [8] and climate change [9], amongst others. Of particular interest in this regard would be the role of these DNNs in image analysis [10] and object detection/segmentation [11].

As a primary fiduciary approach formulated to address these healthcare and environmental issues, the role of the optical microscope cannot be undermined as it aids in the identification of host responses to disease-causing pathogens (e.g. the presence of tumors [12], impact of pathogens on native cellular metabolomics & molecular processes occurring *in vivo* [13], [14], etc), as well as the detection of ecologically-essential microbiota (e.g. diatoms and phytoplankton, such as *Euglena* [15], coupled with soil microbes such as *P. alcaligenes* [16]) which play an essential role in the upkeep of marine and terrestrial ecosystems. Nonetheless, the optical microscope is plagued by a fundamental constraint – its lateral resolution (as it is often utilized) is confined to a minimum distance of 143nm, often described as the Abbe limit [17]. Numerous approaches (both optical and computational) have thus been proposed to circumvent this limitation. The former comprises mainly of optical nanoscopic/super-resolution (SR) techniques [such as SIM [18], STED [19], PALM [20] & STORM [21] (amongst others)], realized through the addition of specialized but *costly* hardware attachments to the optical microscope, while the latter utilizes DNNs (such as ANNA-PALM [22], Deep-STORM^1^ [23], DeepZ [24] or 3D RCAN [25]) in seeking to super-resolve micrographs acquired via conventional imaging techniques (such as widefield epi-fluorescence microscopy) *in silico*.

Despite the ubiquitous availability of DNN frameworks for SR imaging, we have come to realize that a number of the DNNs proposed (such as [22] & [23]) were developed to super-resolve micrographs acquired via traditional widefield *epi-fluorescence microscopy*, where there is a clear distinction between the signal and the background (which is often dark). Such algorithms may thus also perform expectably well for other microscopical imaging modalities such as darkfield microscopy [26] or cross-polarized light microscopy [27], where the background is demarcated by an extinction of optical signals. In contrast, for imaging modalities where there is little discrimination between the background and the signal being resolved, or where artifacts (such as halos or pseudo-relief features) are present in the image due to the imaging technique utilized (as is evident in DIC or PCM), some of these algorithms may not be suitable for super-resolving the input images in this context (as depicted by the implementation of ANNA-PALM [22] in the Results section of the present study). Consequentially, we have sought to develop a novel framework (named O-Net) which we described in our previously published study [28], although the potentiality of further improvements to the O-Net framework were noted to be plausible in its subsequent refinements. In this regard, we now propose an *extension* of the O-Net architecture (which we term Θ-Net) as a viable means of super-resolving micrographs acquired through phase-modulated optical microscopical techniques (namely PCM & DIC microscopy). Θ-Net employs a concatenated architecture of multiple O-Nets (i.e. a computational ‘*string of beads*’), coupled with the transfer learning of features across these different phase-modulated microimaging modalities, so as to exploit learning paradigms of phase-sensitive features resolvable through complementary phase-microscopy techniques. Diagrammatically, the Θ-Net architecture may be represented by Figure 1 as follows:

**Figure 1:**
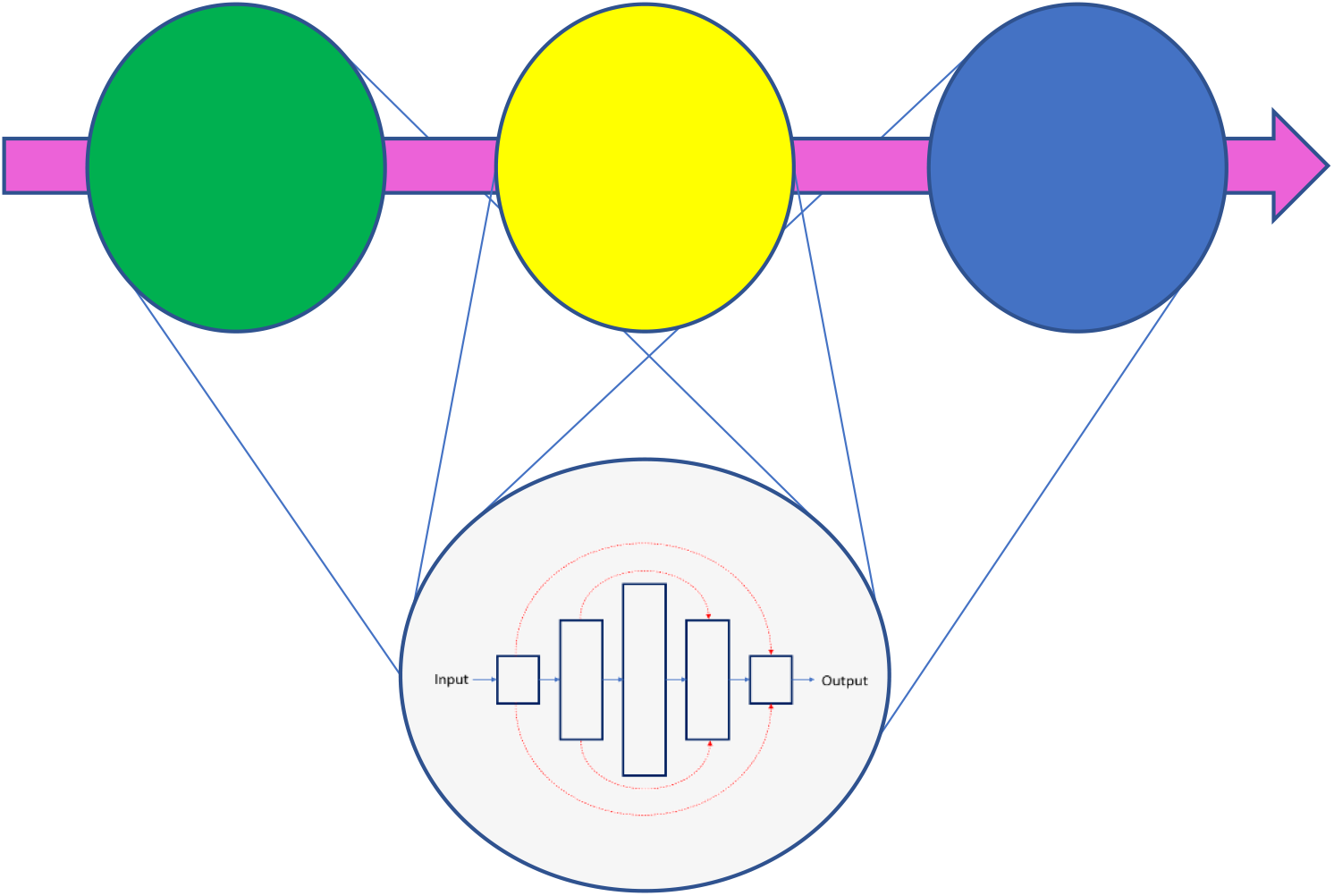
A framework of the Θ-Net architecture as proposed in the current context. Θ-Net adopts a ‘*string of beads*’ methodology of concatenating multiple O-Nets, thereby enhancing the DNN resilience to feature-based variations present in the different samples which it might be trained with. Here (and as is presented in the current study), we employ a 3-node Θ-Net framework for model training & validation.

Testing of Θ-Net was facilitated through attempting to visualize nanoscopic poroids in the striae of the diatom *P. dactylus* var. *dariana* described to be separated by a lateral distance of 72nm – 89nm [29] as verified through SEM (see Figure 2 for details). Here, the capability of Θ-Net in super-resolving PCM & DIC micrographs acquired of this diatom test slide is depicted in the Results section of the present study. In this aspect, we demonstrate that Θ-Net bears significant potential for deployment in label-free optical nanoscopy in the near future, a goal highlighted in [30].

**Figure 2:**
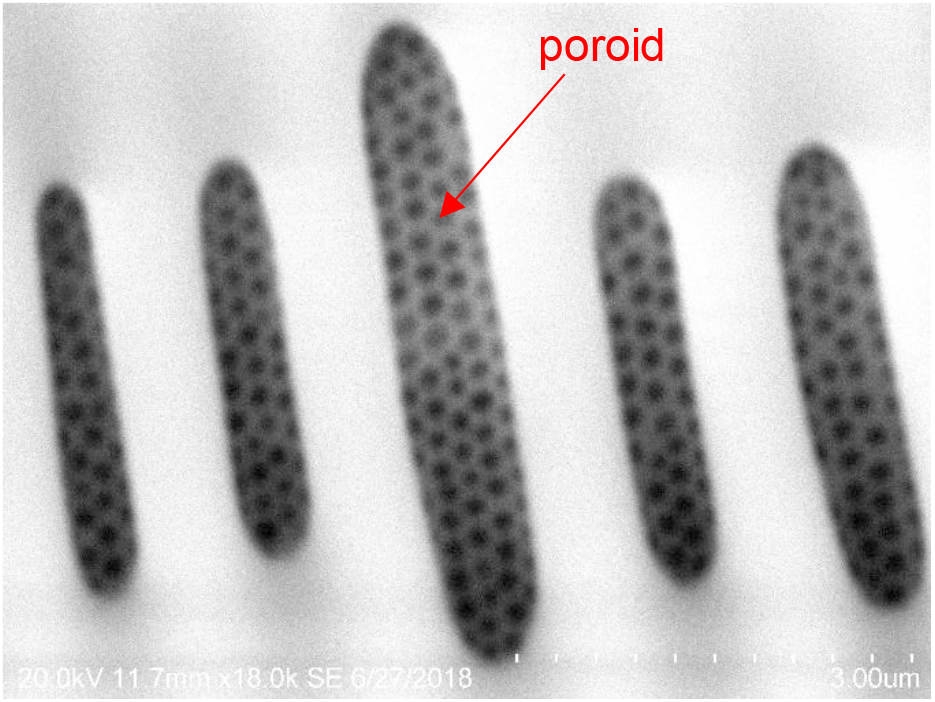
Poroids in the striae of a pennate diatom. Similar poroids are present within the striae of the diatom test sample used in this study (*P. dactylus* var. *dariana*), although the mean inter-poroid spacing for this diatom ranges between 72nm – 89nm, well below the Abbe resolution limit of optical microscopy (typically at ~143nm [17]).

## Materials and Methods

### 1. Data Acquisition & Preparation (Light Microscopy)

The image datasets used for training were similar to those used for training O-Net, as expounded in our previous article at [28]. Generally, commercially available, prepared microscope slides (utilizing samples from plants, animals and microbiota) were used for image acquisition to create the training datasets for the assessed DNNs. The images were acquired through 2 primary imaging modalities – PCM & DIC microscopy – using a Leica N PLAN L 20X/0.4 Corr Ph1 objective (Leica P/N: 506058) [for low-resolution (LR) images] and a Leica HCX PL Fluotar L 40X/0.60 Corr Ph2 objective (Leica P/N: 506203) [for high-resolution (HR) images] installed on a Leica DM4000M microscope, with a CMOS camera (RisingCam® E3ISPM12000KPA, RisingTech) having a pixel size of 1.85μm x 1.85μm and an EK 14 Mot motorized stage (Märzhäuser Wetzlar GmbH & Co. KG) mounted on it. Control of the motorized components of the microscope and camera settings was facilitated through a self-developed stage controller and a desktop UI (also developed in C#). For each sample, similar regions of interest (ROIs) were imaged (under both PCM & DIC microscopies), with the acquired images registered & cropped using MATLAB R2020a (© 1984-2020, The MathWorks, Inc). Where present, shifts detected in the ROIs were mitigated through multi-layer image cropping using Corel PHOTO-PAINT X7 (© 2015 Corel Corporation), prior to splitting into 600px × 600px RGB image tiles in MATLAB R2020a (© 1984-2020, The MathWorks, Inc). These tiles were then conjoined into LR-HR image pairs [the LR images being the **Source** (to be transformed), and the HR images being the **Expected** (*target/ground truth*) images], resulting in a total of 3944 image pairs for each imaging modality employed (i.e. DIC or PCM), before being downscaled to a 256px × 512px format & parsed into NumPy arrays for training the individual nodes of the proposed Θ-Net network in Python 3.8.

For validating the efficacy of the Θ-Net models in image SR, a commercially-available prepared slide of *Pinnularia dactylus* var. *dariana* (*P. dactylus* var. *dariana*) (A. W. F. Schmidt) Cleve 1895 (made at Diatom Shop, Diatom Lab) was imaged separately (through both DIC & PCM microscopies) with a Leica HCX PL Apo 100X/1.4 Oil Ph3 CS objective (Leica P/N: 506211) mounted on the same imaging setup, and the acquired images super-resolved for comparison against a SEM standard. *P. dactylus* var. *dariana* was used as a test sample in this context, as the inter-poroid spacing of 72nm – 89nm [29] is considerably smaller than the computed Abbe diffraction limit of the optical microscope (i.e. 143nm [17]), hence any visible attempt of the tested models to resolve these poroids may be regarded as substantive evidence exemplifying the capabilities of the said models in attaining label-free computational SR microscopy.

### 2. Θ-Net Architecture

The currently-proposed framework (Θ-Net) utilizes a ‘*string of beads*’ architecture, comprising of multiple (in this context, three) O-Nets as described in [28]. The O-Net models utilized for each of the nodes in the Θ-Net scaffold were selected after multiple empirical runs evaluating model architectures of varying depths and trained over a range of epochs (up to a maximum of 320 epochs). Further details pertaining to the individual model architectures are described in the following sub-sections.

#### 2.1 Generalized Structure Underpinning Θ-Net Framework

In the present study, we employed a 7-layer O-Net model trained with the DIC dataset for both the 1^st^ and 2^nd^ nodes, as well as a 7-layer O-Net model trained on *both* the PCM & DIC datasets (the latter of which is *transfer-learnt*) for the 3^rd^ node of the Θ-Net architecture used for super-resolving the input DIC images. For the PCM images, we used a 5-layer O-Net model for the first node, but a 7-layer O-Net model for the 2^nd^ node. For the third node, we use the same transfer-learnt O-Net model as was used for the DIC images. Figure 3 below depicts the general schematic of the Θ-Net framework utilized in this regard:

**Figure 3:**
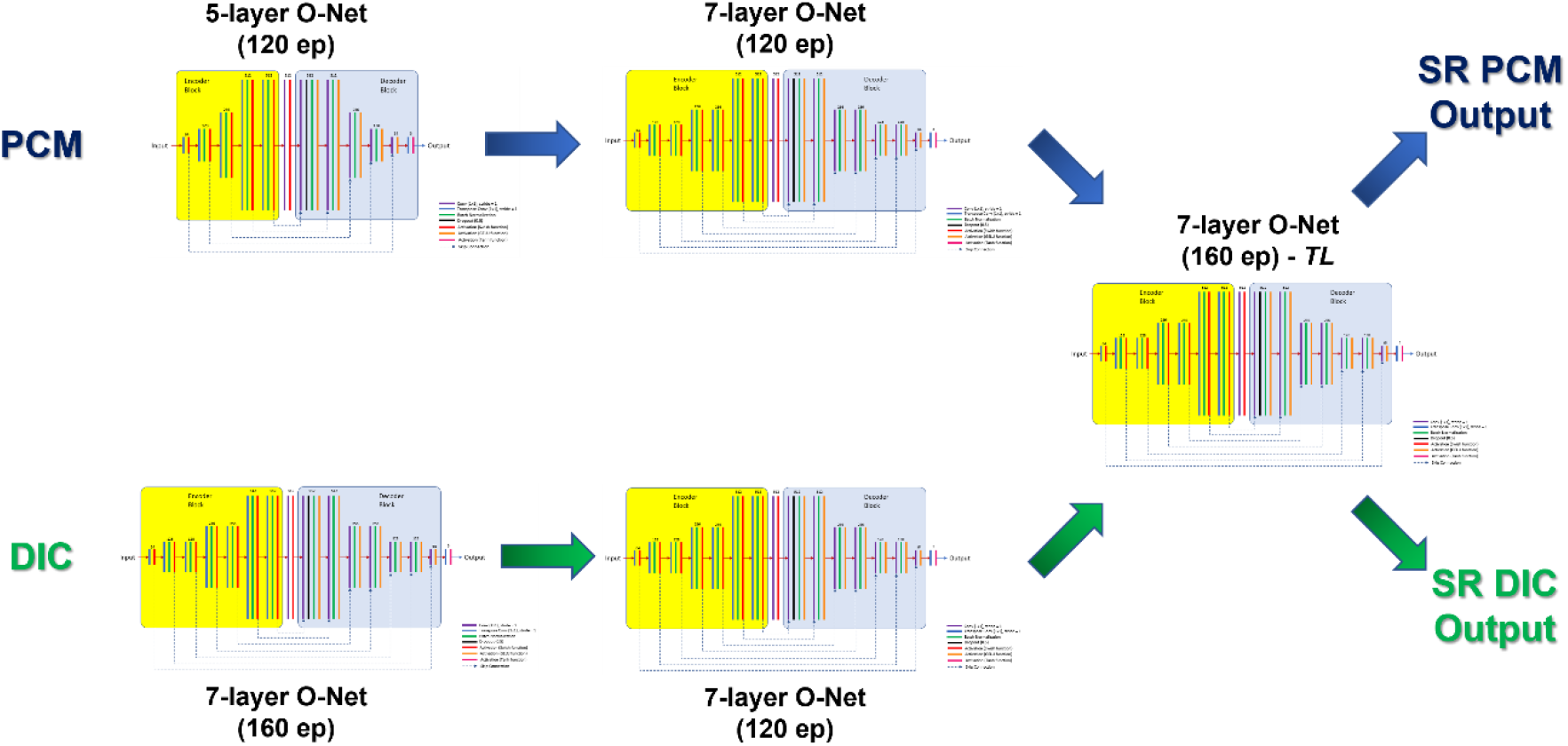
Structure of the 5-layer & 7-layer O-Net architectures utilized as nodes for Θ-Net, as described in the present study. Each of the 3 nodes of Θ-Net (shown in Figure 1 previously) is an O-Net model specifically trained with the input image dataset for the imaging modality which it is intended to be deployed for use in. The exception here refers to the 3^rd^ node utilized in the current Θ-Net framework – the O-Net model here was trained using ***both*** the DIC & PCM image datasets, via a *transfer learning* approach. As with the traditional O-Net architecture (described in [28]), skip-connections (*concatenations*) are used to join layers in the encoder block (consisting of transposed convolution operations) with their corresponding conjugates in the decoder block (comprising of convolution operations).

At this juncture, it would be essential to highlight that the individual O-Net nodes within a Θ-Net chain are founded on the Pix2Pix GAN architecture, as discussed in [28] & [31]. Similarly, Swish [32] & GELU [33] activation functions were also incorporated into the encoder & decoder blocks respectively (the mathematical definitions of these functions being included in the accompanying Supplement for the interested reader). The images generated by the Θ-Net models were then analyzed in MATLAB R2022b (© 1984-2022, The MathWorks, Inc) using standard image quality metrics, such as the peak signal-to-noise ratio (PSNR), signal-to-noise ratio (SNR), image mean-squared error (IMSE) & structural similarity index (SSIM). The findings gleaned from this assessment are henceforth depicted in the Results section of this study.

#### 2.2 Cross-domain learning

An interesting aspect of the present study refers to the employment of *cross-domain learning* [34] across 2 commonly utilized phase-modulated optical microscopy techniques (i.e. PCM & DIC microscopies). Here, we demonstrate that it is possible to extricate information gleaned from images acquired under each of these techniques to *enhance* the overall Θ-Net models performance in computational nanoscopy (as exemplified in the Results section of this study). This may potentially be attributable to **both** DIC & PCM translating *optical path length* (OPL) variations in the sample into amplitude (image brightness/intensity) differences, albeit via different routes (PCM translates differences in OPL *magnitude* into brightness variations, while DIC converts OPL *gradient* differences into 3-D relief effects [35]). In this respect, we seek to utilize *transfer learning* as a vehicle to facilitate cross-domain learning between these 2 microscopical imaging modalities, thereby spelling benefits for the Θ-Net-generated SR images to incorporate the ‘best of both worlds’, while being potentially independent of the limitations faced when employing each technique individually (e.g. birefringence artifacts encountered in DIC, as well as a lack of lateral & axial resolution coupled with an unsuitability for specimens having high phase shifts for PCM imaging [35]).

#### 2.3 Comparative Networks for SR Image Analysis

Separately, we have also attempted to include models derived from other industry-leading DNN frameworks (such as ANNA-PALM [22] & 3D RCAN [25], the latter of which is employed for *in silico* nanoscopy in the Aivia deep-learning suite [36]), as a comparative performance gauge of the proposed Θ-Net models. It would be apt to emphasize (as was also highlighted in the Introduction section earlier) that these frameworks have been *specifically* adapted for super-resolving *widefield epi-fluorescence monochrome/grayscale* microscopical images (3D RCAN [25] uses grayscale image Z-stacks for generating the SR image), hence a somewhat *close* (albeit not very similar) comparative assay in this respect would require splitting the current 2D RGB image into a 3-channel grayscale image stack, which is then simulated as a Z-stack for training & validating the 3D RCAN [25]-based models. For ANNA-PALM [22] validation, we utilized the ImageJ A-Net plugin downloaded from [37] and the supplied models trained for super-resolving microtubules within ImageJ, as the structures we wanted to super-resolve (i.e. the meshwork between the poroids in the striae) may be interpreted to resemble (in part) the filamentous strands characteristic of microtubules. In addition, models developed on a third comparative framework (termed BSRGAN [38]) were also included in the present study, for verification purposes.

### 3. Image Denoising

In addition to image SR, we have also attempted to evaluate the capability of the proposed Θ-Net models for *image denoising*, despite the models not being trained specifically for this purpose. To do this, salt-and-pepper noise was synthetically introduced into the validation images using MATLAB R2022b (© 1984-2022, The MathWorks, Inc) and the assayed Θ-Net models assessed on their propensity to denoise the noise-infused images.

The codes and models used to generate and evaluate the images presented in this study are available for download in the accompanying Supplement.

## Results

The models developed using the presently-assayed Θ-Net architecture were contrasted against our previous O-Net models [28], as well as other state-of-the-art *in silico* SR architectures, such as 3D RCAN [25], BSRGAN [38] & ANNA-PALM [22] for each of the DIC & PCM imaging modalities. The generated images obtained from each of these models are depicted in the subsequent figure:

### 1. DIC Imaging

From Figure 4, it may be clearly discerned that images generated via the Θ-Net-trained models exhibit an enhanced contrast, and consequently, an increased resolution of details and features as compared to those generated from the O-Net-trained models. Nonetheless, pseudo-relief depictions of SR features are also apparent in the Θ-Net-generated images, indicative of the resemblance borne to the **Source** (input) images, rather than the output (**GT**) images. This may exemplify a need for further training of the Θ-Net models, potentially via additional nodes/*beads* appended to the existing ‘*string (of beads)*’ characteristic of the Θ-Net architecture, although network hallucination (a prominent flaw in DNN-generated images as highlighted in [39]) is not clearly evident in both the O-Net and the Θ-Net-generated images.

**Figure 4:**
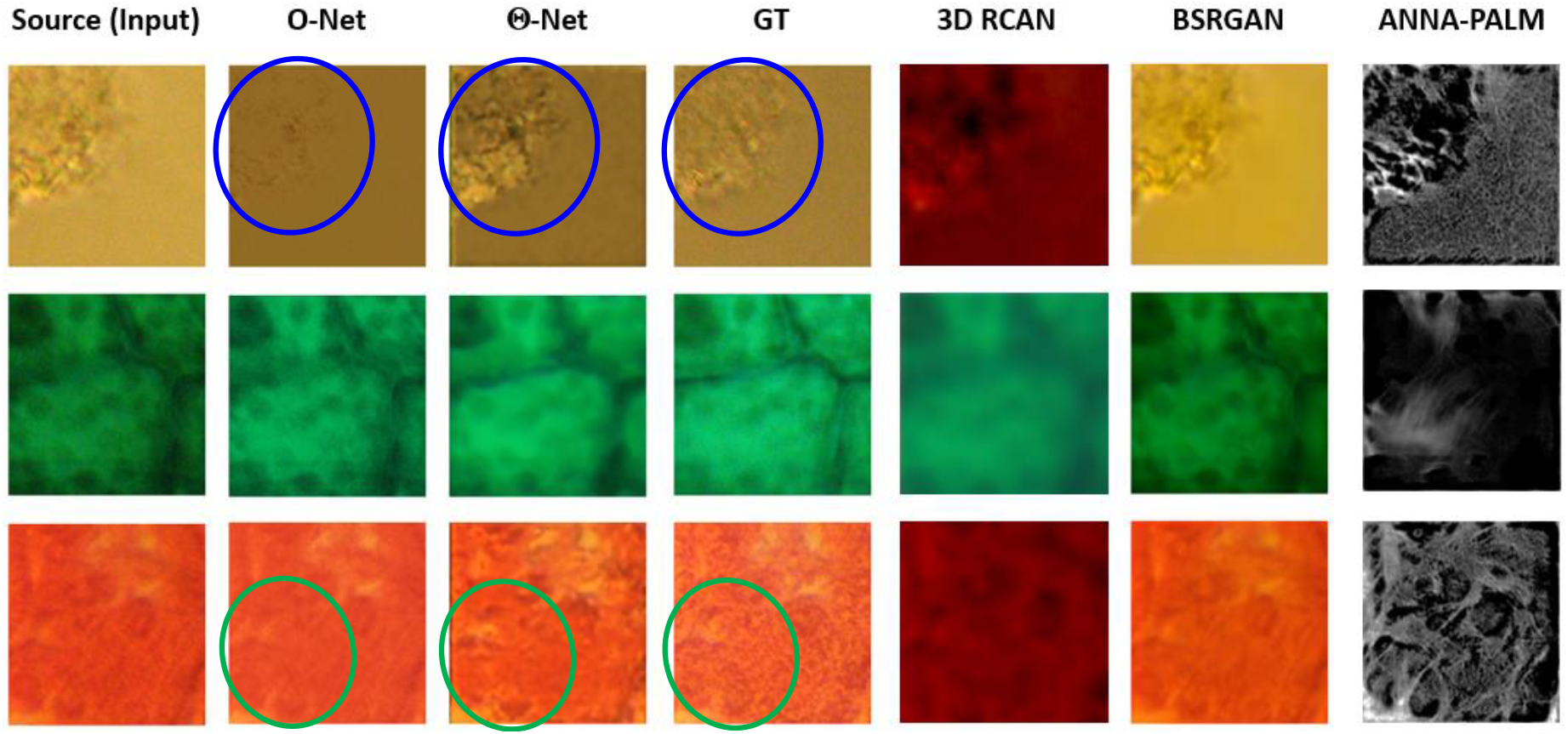
Image SR obtained through several models, including those developed using O-Net [28] & Θ-Net. The O-Net model was trained over 101 epochs, while the Θ-Net model assimilated O-Net models trained over 160 epochs (for the 1^st^ node), a 120-epoch-trained O-Net model (for the 2^nd^ node) and a 160-epoch-trained & transfer-learnt O-Net model (for the 3^rd^ node). Notice the closer resemblance of the Θ-Net-generated image to the **GT** (Ground Truth) image, as compared to the O-Net models (highlighted within the blue and green ellipses). *N.B.: The **Source** image (input) was acquired via the 20X/0.40 Ph1 objective, while the **GT** (Ground Truth) image was obtained using a 40X/0.60 Ph2 objective. Images generated through models founded on other frameworks (namely 3D RCAN [25], BSRGAN [38] & ANNA-PALM [22]) were also included for comparison purposes (the ANNA-PALM [22] model developed for super-resolving grayscale photomicrographs of microtubules was utilized as an extension within ImageJ, while the 3D RCAN [25] model was trained over 250 epochs with 1972 steps per epoch & 2 residual groups).

Approaching from a computational stand-point, we have indicated the plots of the respective loss functions, namely the discriminator losses on real samples (dR) & generated samples (dG), as well as the generator (g) loss for the training runs in the various nodes of the Θ-Net models in Figure 5 below:

**Figure 5:**
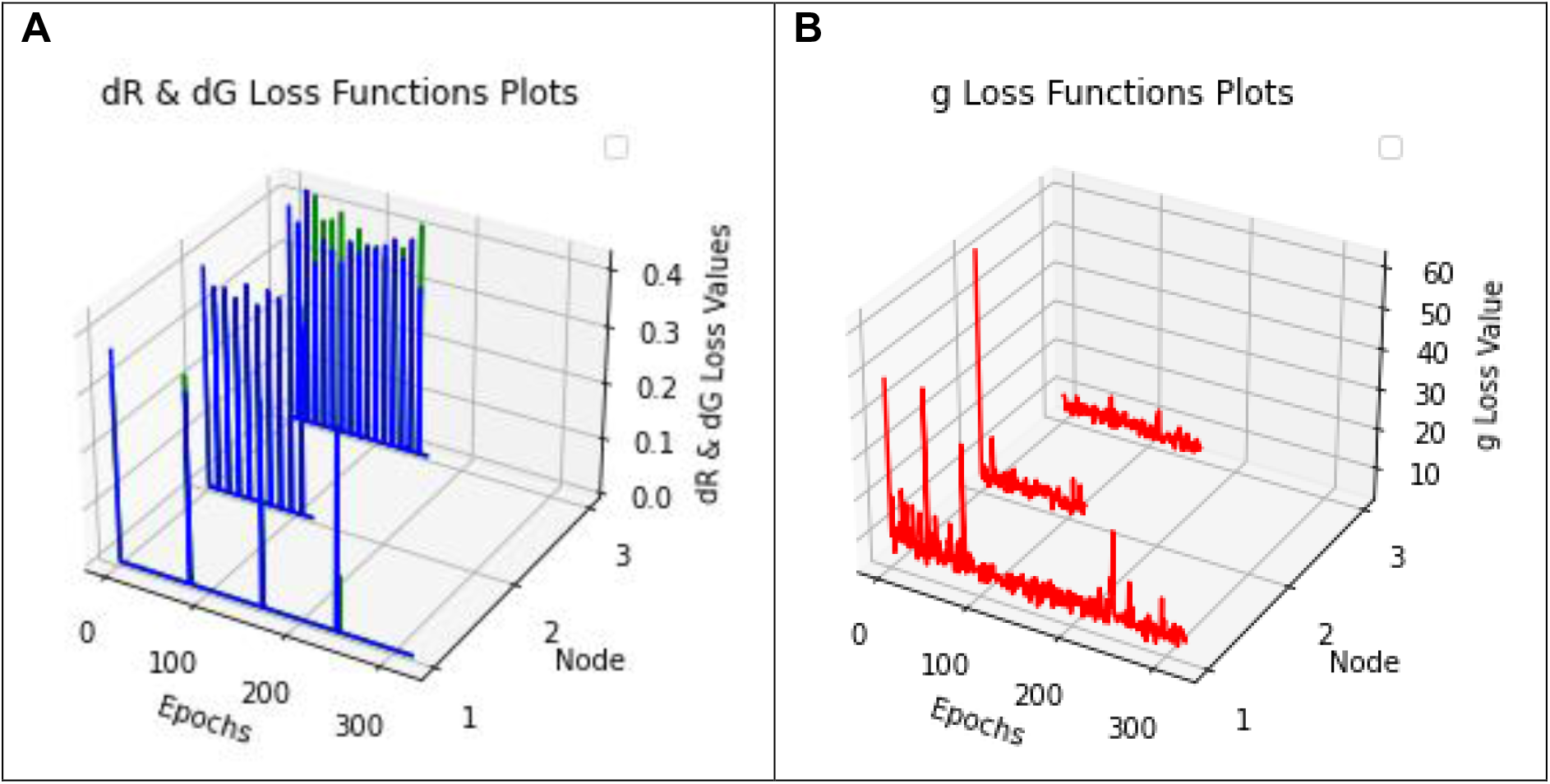
Loss functions for DIC micrographs imaged under the Θ-Net architecture. **A** Discriminator losses on *both* the real & generated samples, represented by dR (green line) & dG (blue line) losses respectively, as well as **B** the generator (g) loss plotted as a red line. The spikes observed in the dR and dG loss function plots (**A**) demonstrate the commencement of a subsequent training run, implicating the selection of a new random seed for the DNN training.

Similarly, analysis of the images acquired via PCM revealed the following observations (detailed in the subsequent sections):

### 2. PCM Imaging

Analyses of the images portrayed in Figure 6 depict a similar trend to that observed for Figure 4 (i.e. that Θ-Net-generated images more closely resemble the ground truth images as compared to O-Net-produced images), favoring the *string of beads architecture* hypothesis proposed in the present study to attain image SR in computational nanoscopy. Nonetheless, as with Figure 4 previously, there still exists room for improvement when comparing the effective resolution of the Θ-Net-generated images with the ground truth images, implying that more nodes/*beads* could be added to the *string of beads* structure characteristic of Θ-Net in a bid to improve image SR. As with the Θ-Net models utilized for super-resolving DIC images earlier, we have sought to plot the respective loss functions (dR, dG & g loss) for the various Θ-Net models, as shown in Figure 7 below. Here too, a similar legend is being used for the plots – blue is indicative of dG loss, green for dR loss & red for g loss.

**Figure 6:**
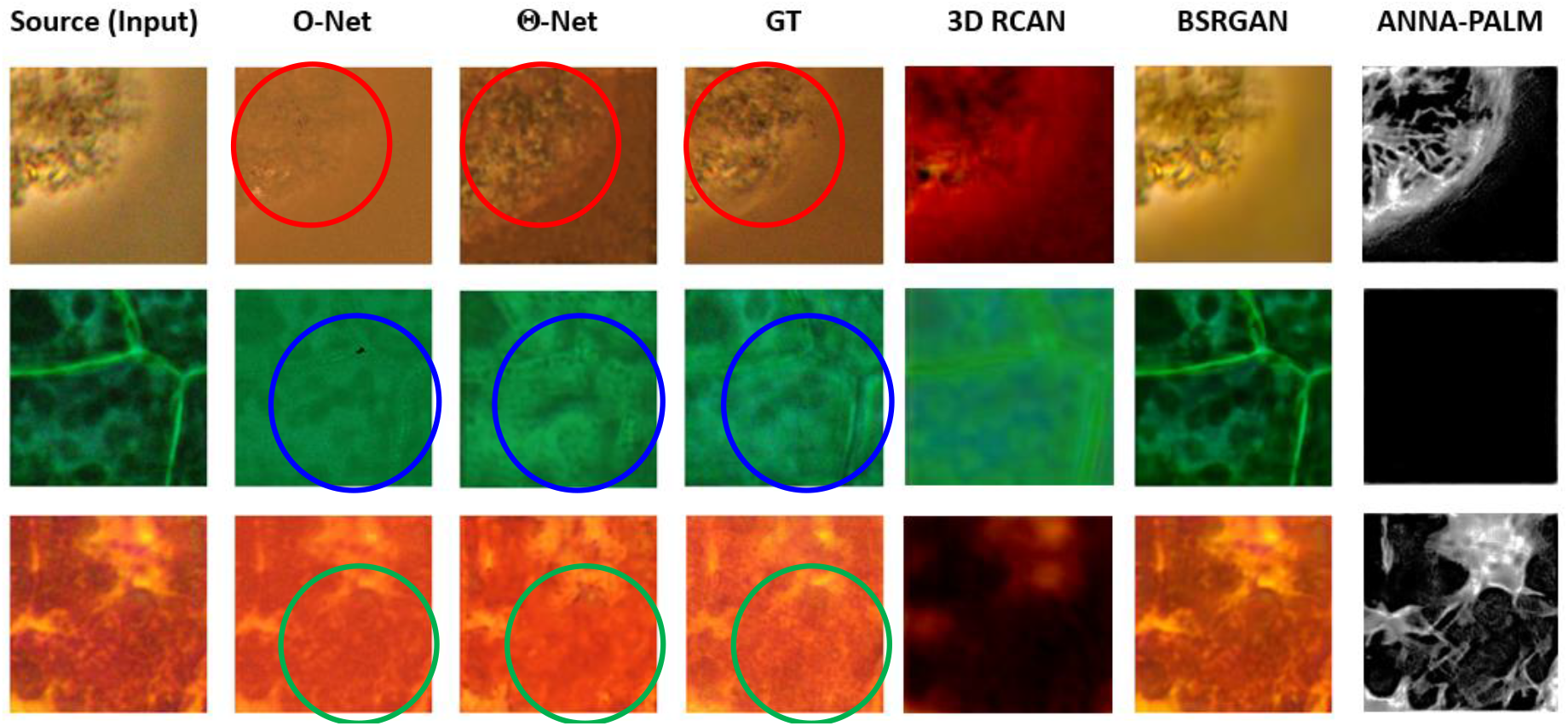
A comparison of the images obtained through utilizing models developed on the O-Net and Θ-Net architectures using PCM micrographs. Here, the O-Net model was trained over 120 epochs, while the Θ-Net framework incorporated the said O-Net model (in its 1^st^ node), a 120-epoch-trained O-Net model (for the 2^nd^ node) and a 160-epoch-trained & transfer-learnt O-Net model (for the 3^rd^ node), which is identical to that used for the DIC micrographs in Figure 4 previously. Also (as enunciated for the images in Figure 4), the **Source** (Input) image was acquired via the 20X/0.40 Ph1 objective, while the **GT** (Ground Truth) image was obtained with a 40X/0.60 Ph2 objective). As with the Θ-Net-generated DIC micrographs in Figure 4 previously, the regions encircled within the red, blue and green ellipses of the Θ-Net-generated images depict a closer correlation with their corresponding **GT** images (as compared to their O-Net counterparts), supporting the presently-proposed Θ-Net framework as a viable improvement over O-Net, for producing computational models to facilitate *in silico* SR. Here too (as with Figure 4 previously), the ANNA-PALM [22] model was utilized as an ImageJ extension for microtubule SR, while the 3D RCAN [25] model was trained over 250 epochs with 1972 steps per epoch & 2 residual groups.

**Figure 7:**
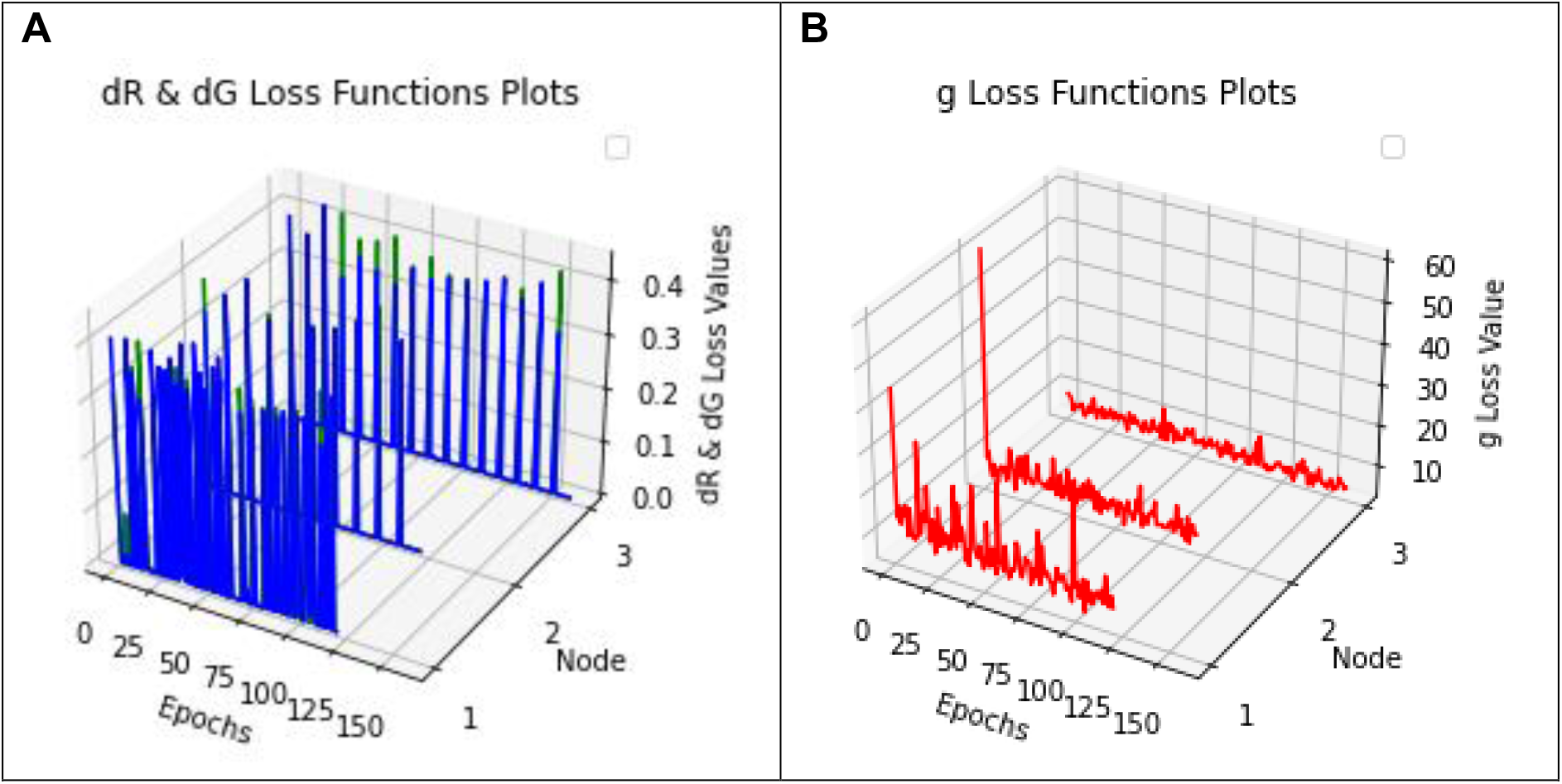
Loss function plots for Θ-Net models trained on PCM micrographs. **A** denotes the discriminator losses for real (dR) & generated (dG) samples, indicated by green & blue lines respectively, while **B** represents the generator (g) loss (as red lines). As with Figure 4 previously, spikes present in the plots of the dR & dG loss functions indicate the start of a new training run (using the existing Python script), potentially implying the selection of a new random seed at the start of each run (model training in the present context was conducted intermittently).

Here, it would be vital to mention that when evaluating the performance of a generative adversarial network (GAN), such as that of Θ-Net, there is ***no*** objective loss function which can be used to do this [40], hence we have chosen to utilize the discriminator losses on the real and generated samples (as well as the generator loss) as key determinants for assessing Θ-Net performance. We have done this for the Θ-Net models trained on *both* the DIC & PCM datasets (as depicted in Figure 4 & 7 respectively). The formulae underlying these losses are indicated in the Discussion Section of this study.

### 3. Computation of Global & Local Image Metrics

Further evaluation of the global & local image metrics (for specific ROIs in a subset of the validation images as shown in Figures 4 & 7 previously) was subsequently conducted, and the results indicated in Figure 8:

**Figure 8:**
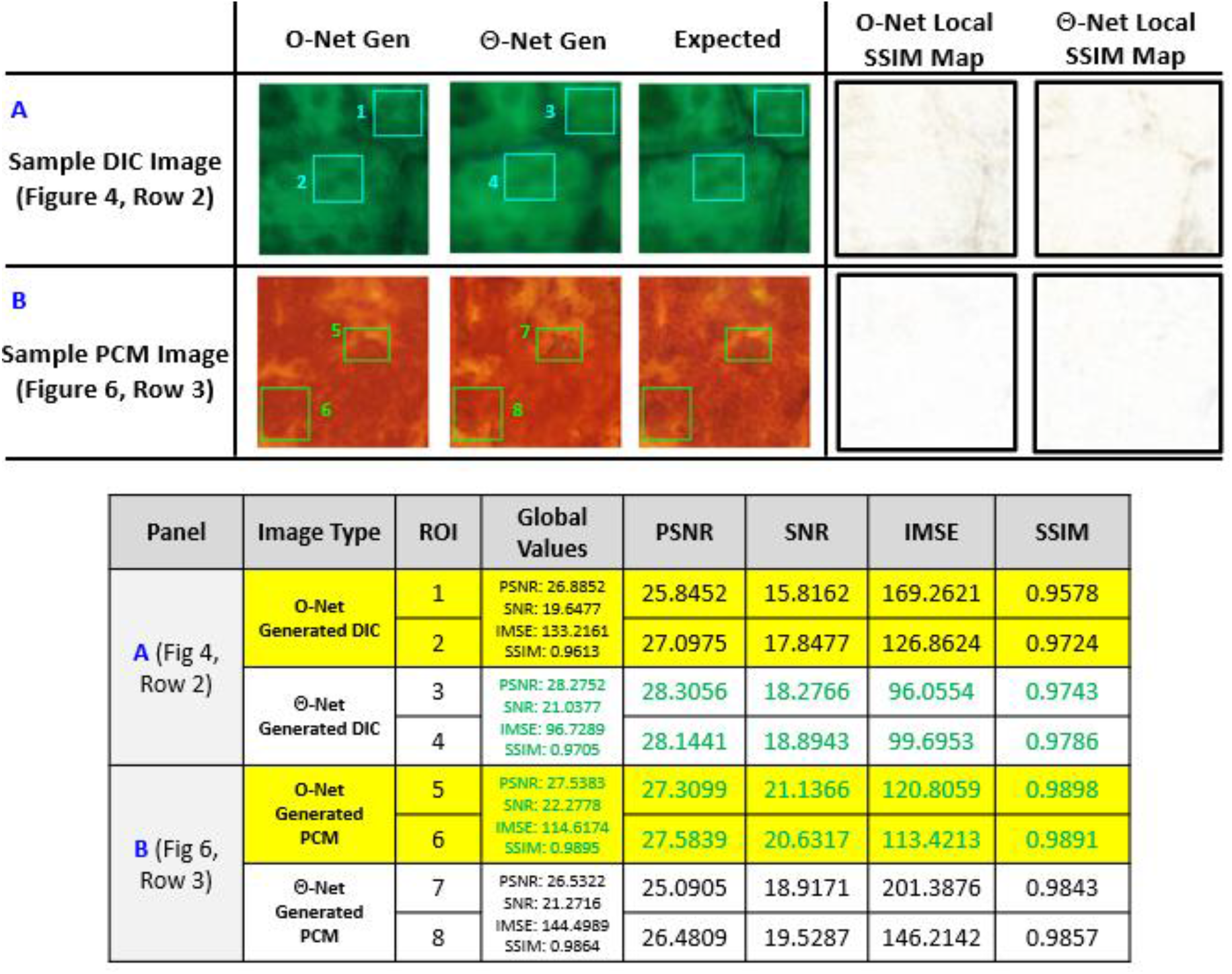
Sample O-Net & Θ-Net-generated images extricated from **A** DIC micrographs (Figure 4, Row 2) and **B** PCM micrographs (Figure 6, Row 3). In this instance, we have intentionally selected images where it was highly likely that Θ-Net *underperformed*, as a means of exercising stringency when evaluating potential mismatches between Θ-Net & ground truth (**Expected**) images. Here, we observe that the local SSIM Maps for both O-Net and Θ-Net appear to be generally white, which is indicative of a high correlation between the respective DNN-generated images and the ground truth (**Expected**) images [the local SSIM maps are based on individual pixel mismatches within an 11-by-11 neighborhood [41] & range from black (0) to white (1)]. From the table, it may be observed that Θ-Net generally surpasses O-Net when super-resolving DIC micrographs, while the reverse holds true for PCM images (at least for the assayed images in this instance). Nonetheless (for the PCM images), the image metrics (such as PSNR and SSIM) for Θ-Net seem to closely approach those of O-Net (differing by <1% for SSIM scores but a slightly higher margin of <9% for PSNR scores), implying that Θ-Net models might be more susceptible to local variations in individual pixel values and seeking to convert these into discernible features (which may consequentially result in an imposed ‘penalty’ and reduced SSIM scores, as these metrics may interpret these features as noise). Further evidence of this is provided through the IMSE metric, which clearly shows a marked increase for the Θ-Net-generated images (when compared with those from O-Net) for the PCM images (Panel **B**).

From the findings in Figure 8, we may deduce that Θ-Net generally performs relatively well in super-resolving phase-modulated micrographs (the figures chosen in this context were selected from images where a lower performance of Θ-Net was expected). Despite this fact, regions where Θ-Net was noted to have under-performed (when compared to O-Net) were prominent in the PCM image dataset, although further inspection revealed that this reduced performance may be attributed to Θ-Net seeking to super-resolve fine structural details within the source image, resulting in lowered PSNR scores (since this metric is often used to penalize noise, while the latter may be reminiscent of *pseudo-noise* as discussed in [28]). In this context, we would thus need to evaluate the performance of Θ-Net from the perspective of other metrics as well (such as SSIM), since these provide a holistic representation of Θ-Net as a DNN framework for *in silico* SR microscopy. Judging from this angle, we notice that the SSIM scores of Θ-Net-generated images closely approach those of O-Net (differing within 1%), implying that there is very little difference between the quality of Θ-Net and O-Net-generated images.

A subsequent validation was performed to evaluate the *true* efficacy of the proposed networks in super-resolving both PCM & DIC images of a specimen **never** trained before with either network. The test specimen was that of a pennate diatom (*P. dactylus var. dariana*) with a consequent aim to resolve the sub-microscopic poroids in the diatom striae. The results of this validation study are presented in the following Figure 9:

**Figure 9:**
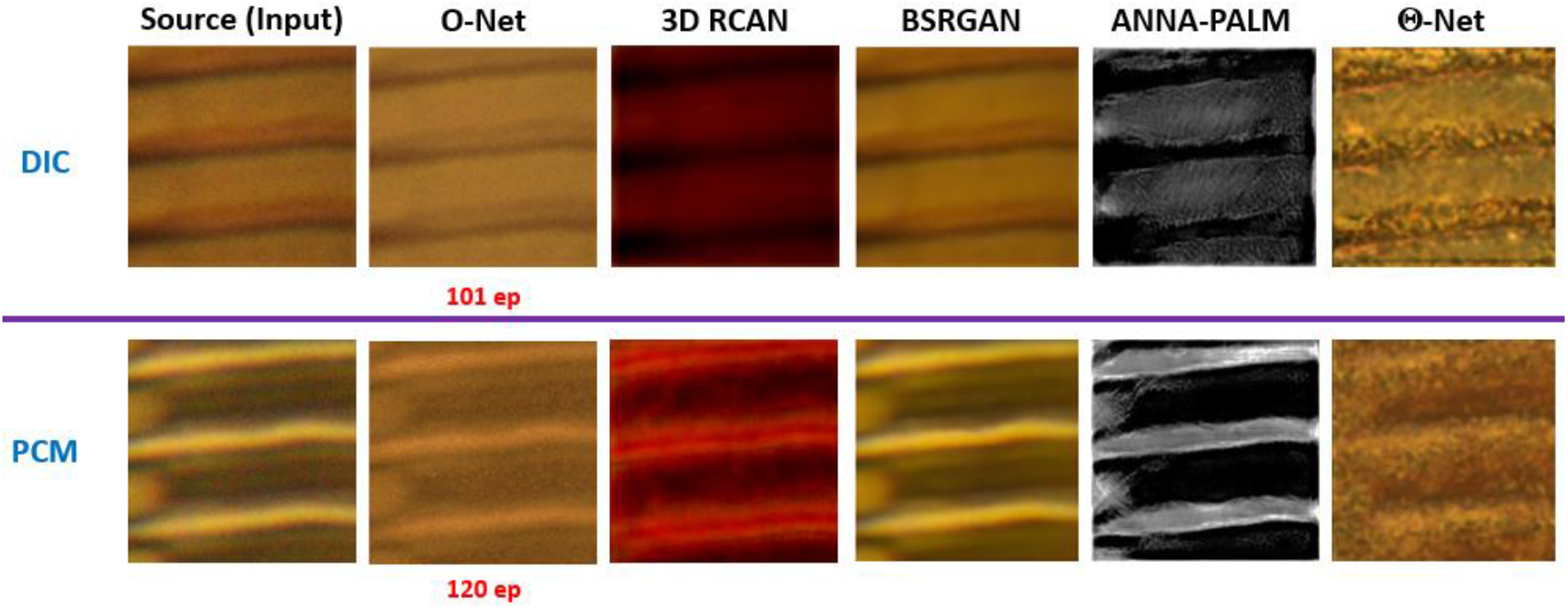
DIC & PCM micrographs of *P. dactylus* var *dariana* acquired using a 100X/1.4 Oil Ph3 objective (Leica P/N: 506211) [**Source (Input)** images] and super-resolved through application of models adopting the O-Net & Θ-Net architectures, for both PCM & DIC micrographs (3D RCAN [25], BSRGAN [38] & ANNA-PALM [22] are also included for comparison purposes in the present context). Here, it is obvious that the surface of the diatom striae appear to be significantly more granular for Θ-Net-generated images than O-Net-generated images, an observation which closely correlates with that of the SEM photomicrograph of *P. dactylus* var *dariana* as shown in Figure 2 previously.

Closer inspection of Figure 9 (with particular attention to the images generated by Θ-Net & O-Net) reveals the presence of bright specks within the darker meshwork of the diatom striae – a feature postulated to be the poroids of *P. dactylus* var *dariana* being characterized by [29] to have been separated by a mean distance of 80.5nm ± 8.5nm, thus lying well below the effective Abbe resolution of the optical microscope (of 143nm) [17]. In this respect, we may surmise that Θ-Net models do indeed perform well as DNN-mediated regulators for phase-modulated nanoscopical applications, even in the context of blind *in silico* SR (in the absence of *a priori* information on the optical PSF/OTF of the system).

### 4. Image Denoising

To evaluate the efficacy of the proposed models in *denoising* input micrographs (even though the assayed models were **not** specifically trained for this purpose), we have infused an artificial representation of noise into the source images via a salt-and-pepper noise algorithm in MATLAB, with the image denoising consequently performed using the same trained models in Python. The results of this trial [including the corresponding source & ground truth (**Expected**) images] are depicted in Figure 10 as follows:

**Figure 10:**
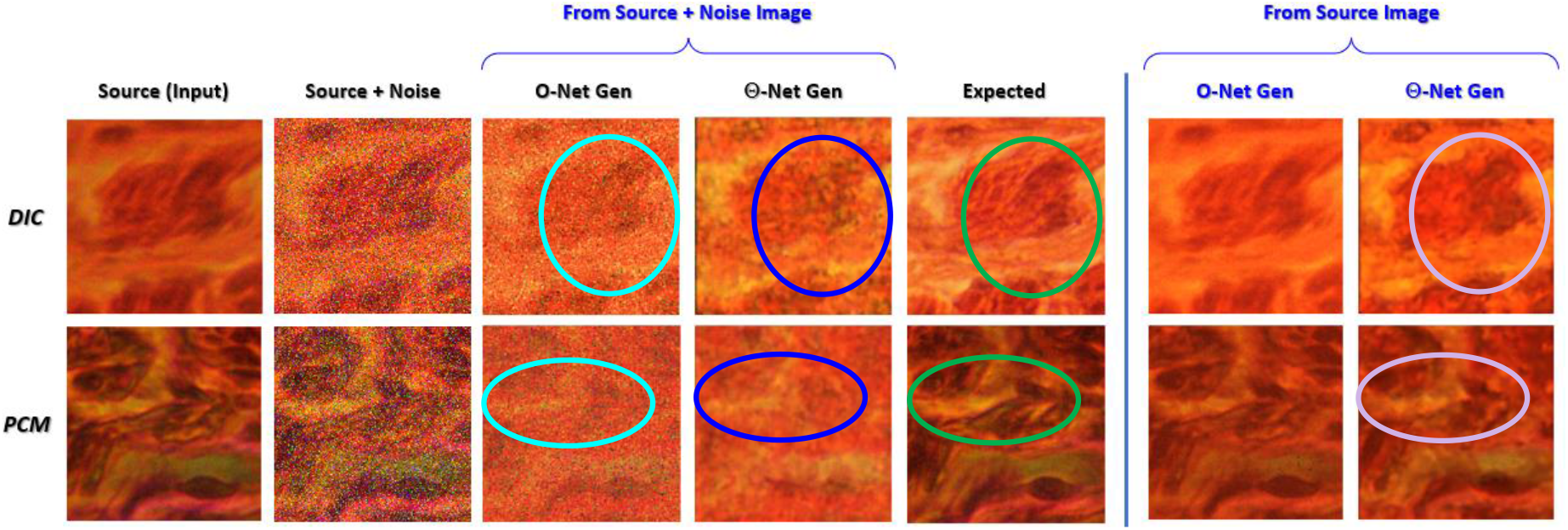
DIC and PCM photomicrographs infused with salt-and-pepper noise (noise density: 20%) which are labelled **Source + Noise**, and their corresponding ‘denoised’ images, as well as the ground truth (**Expected**) images utilized here. Here, we notice that Θ-Net models are particularly sensitive to these noisy pixels, misrepresenting them as ‘features’ (artifacts) within the generated images (as indicated within the blue ellipses) when compared with the ground truth (**Expected**) images (demarcated within green ellipses), thereby potentially suggesting the emergence of *network hallucination* in this context. This phenomenon is not as evident in the O-Net-generated images though (indicated within cyan ellipses), implying a greater resilience of O-Net based models to noise variations of this type. It is thus evident that Θ-Net (though highly viable & desirable for *in silico* SR as portrayed in the previous Figures 4, 6, 8 & 9) does ***not*** perform well in the presence of salt-and-pepper noise, implying a need to *denoise* the Source (input) images *separately*, prior to feeding them into the Θ-Net models for further downstream processing (i.e. image SR). This deduction is further corroborated by the absence of such artifacts when evaluating the Θ-Net-generated images in the absence of input noise (depicted within the violet ellipses). Nonetheless, it would be prudent to mention at this juncture that the Θ-Net models perform very well in effectively removing the noisy pixels in the image (unlike their O-Net counterparts, where the noisy pixels are still present).

### 5. Computational Complexity & Load

In addition to assessing the quality of SR images generated from each of the assayed architectures (in particular Θ-Net), we have also sought to compare the execution times for the afore-mentioned models, both as a means of quantifying their computational complexity and to determine the viable temporal resolution attainable by each of the models. The results following this analysis are presented in Table 1 below:

**Table 1:** Execution times (in seconds) indicated for each of the models trained under the assayed frameworks (i.e. O-Net & Θ-Net) in the present study. As Θ-Net is comprised of 3 nodes (each of which is an O-Net model), implementation of Θ-Net for image post-processing is expected to take ~thrice as long as that of O-Net, although this is a non-linear relationship, being very much dependent on GPU capabilities.

| Architecture | CPU | GPU | RAM (GB) | Storage | Average Execution Time |
| --- | --- | --- | --- | --- | --- |
| **O-Net** | Intel® Core™ i5-7200U CPU | NVIDIA GeForce® 920MX | 20 GB | 1 TB SSD | 11.132 s (DIC) / 11.463 s (PCM) |
|  | 2 x Intel® Xeon® Platinum 8170 (56C/112T) | NVIDIA Tesla K80 | 128 GB | 500 GB SSD | 2.437 s (DIC) / 2.132 s (PCM) |
|  | Intel® Core™ i9-10920X | NVIDIA GeForce RTX™ 3090 | 128 GB | 512 GB SSD | 0.422 s (DIC) / 0.297 s (PCM) |
| **Θ-Net** | Intel® Core™ i5-7200U CPU | NVIDIA GeForce® 920MX | 20 GB | 1 TB SSD | 51.756 s (DIC) / 40.291 s (PCM) |
|  | 2 x Intel® Xeon® Platinum 8170 (56C/112T) | NVIDIA Tesla K80 | 128 GB | 500 GB SSD | 14.89 s (DIC) / 19.613 s (PCM) |
|  | Intel® Core™ i9-10920X | NVIDIA GeForce RTX™ 3090 | 128 GB | 512 GB SSD | 0.593 s (DIC) / 0.703 s (PCM) |

From the findings tabulated in Table 1, we may observe that the average execution times of Θ-Net (a DNN architecture comprised of 3 O-Net nodes strung together in the present context) expectably exceeds that of O-Net, although this increase does not appear to be proportional to the node-count of the Θ-Net framework used. Instead, the execution times seem to be highly dependent on the type of GPU used, ranging from a maximum multiplier of ~9.2X (for the NVIDIA Tesla K80 GPU system) to a low of ~1.4X (for the NVIDIA RTX 3090 GPU system). As GPUs are known to excel in parallel computationally intensive tasks, this might suggest that the individual O-Net nodes are being executed in a parallel (rather than a sequential) fashion.

## Discussion

The results depicted in this study exhibit the significant potential of Θ-Net in attaining computational phase-modulated nanoscopy. Here, it may be observed that Θ-Net (as with O-Net in [28]) can super-resolve both DIC & PCM micrographs while avoiding the formation of potential artifacts characteristic of these imaging modalities (namely the pseudo-relief effects DIC microscopy [42] & the halo/shading-off effects of PCM [43]). In this regard, we surmise that the Θ-Net-based models likely do this (i.e. image SR) via a mapping function to reduce the PSF/OTF of the optical system (akin to that for O-Net as described in [28]), resulting in a hypothetical PSF which (when convolved with the ground truth representation) produces a SR image of the specimen in question (further details on this are expounded in [28] for the interested reader). However, Θ-Net does this through *repeated functional mapping* of the acquired/ generated PSF-convolved input image, which may be generally described by Equations 1 & 2 below: **<u>Learning Phase</u>**

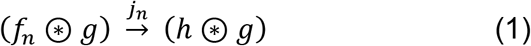

where *f*_1_ refers to the PSF of the optical system when using the 20X/0.4 Ph1 objective, *g* is the ground truth image of the specimen and *h* is the PSF of the optical system (when using the 40X/0.6 Ph2 objective). Here, *f*_n+1_ refers to the *generated* ‘PSF’ of the image when (*f*_n_ ⊛ *g*) is mapped under *j*_n_. Like O-Net [28], the Θ-Net models thus attempt to learn the mapping function *j*_n_ across *n* nodes (for which Θ-Net is defined), and (upon sufficiently learning this) deploys *j*_n_ for image SR as follows: **<u>Mapping Phase</u>**

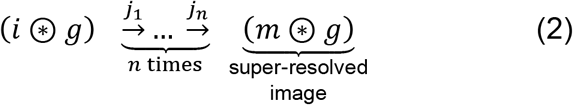

where *i* refers to the PSF of the optical system when using the 100X/1.4 Ph3 objective and *m* refers to the *computed hypothetical* PSF of the optical system for the SR image. Nonetheless, a key aspect of our present study remains – i.e. to highlight the superiority of models adopting the Θ-Net framework over their predecessors employing the O-Net framework (as described in [28]) for *in silico* label-free SR microscopy – evidence for this being presented in Figures 4 & 6 respectively. As previously enunciated, Θ-Net (in the present study) employs a *triple*-node architecture (each node being an O-Net) with a specialized *transfer-learnt* O-Net model for the last node (the said node being trained on *both* DIC & PCM datasets). Here, the transfer learning process clearly aids in enhancing the SR capabilities of the Θ-Net models, putatively by allowing a *transfer* of acquired phase information (manifested as amplitude variations) across 2 principally-similar (yet methodologically-different) imaging modalities (i.e. DIC & PCM) to augment the trained models’ abilities in detecting and translating these phase variations to the feature space. Empirical validation of this statement may be derived from Figure 9, where the granular surface of the striae of the pennate diatom *P. dactylus var. dariana* (coupled with its sub-microscopic poroids) become distinctly visible upon processing with the Θ-Net models (as compared to the O-Net models), closely resembling the SEM gold standard micrograph as depicted in Figure 2. The poroids (though) may be putatively identified as bright pores within the slightly darker mesh of the striae – an effect evocative of light transmission through the poroids in diascopic imaging modalities, such as the ones utilized in the present context.

From a computational perspective, execution times of Θ-Net seem to be viable as well, generally subceding 1s in current advanced GPU systems, making it viable where processes having a temporal resolution exceeding 1s are generally encountered. Moreover, the loss function plots of the assayed Θ-Net models (for both DIC and PCM) suggest that the models have been optimally trained, with the discriminator losses for both real & generated samples (indicated by dR & dG respectively) generally approaching 0 for all three nodes. The equations underpinning the computed losses [dR, dG & the generator loss (g)] may be mathematically defined as follows:

For the discriminator losses (i.e. both dR & dG), the **binary cross-entropy** loss (or log loss) ε was utilized, which may be expressed as below (from [28] and [44]):

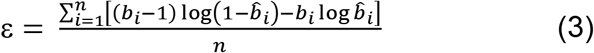

where *b_i_* is the label and 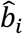 is the probability of *b*_i_ = 1 (derived from [44] & [45], and described in [28]).

In contrast, for computing the generator (g) loss, both ε *and* the **mean absolute error** (MAE) / l_1_ loss were used, the latter being computed as follows(from [28] and [46]):

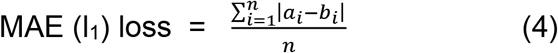

where *n* is the number of pixels in the image, *a_i_* & *b_i_* refer to the target and the estimated values of the assayed parameter [e.g. pixel RGB (or HSL) intensities at pixel *i* respectively]. Compounded together, the overall generator loss G may thus be expressed by the following equation (from [28]):

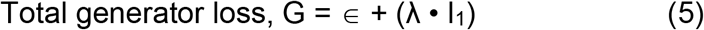

where λ = 100 [47]. ε and MAE were utilized as viable loss functions for evaluating model training in this respect as they provide viable indications on how closely the generated images resembled the ground truth (target) image under differing representations (MAE evaluates this from a linear perspective, while ε adopts a logarithmic standpoint).

On a separate note, empirical verification of the image denoising capabilities of the Θ-Net models suggests that these models are particularly prone to salt-and-pepper noise present in the images, confounding this with sub-microscopic features which require a modulation transfer function (MTF) greater than that afforded by the microscope’s optical train [48]. Evidence for this may be drawn from Figure 10, where some features encircled within the blue ellipse of the DIC micrograph depict an attempt by the said model to translate some of the noise into ‘super-resolved features’ (a characteristic generally absent in the traditional O-Net-based models). This finding thus portrays the need for the Θ-Net user to *denoise* the input images separately, prior to super-resolving them via the proposed models. Despite this fact (and as previously already highlighted in the caption for Figure 10), the Θ-Net models perform significantly better at removing the noisy pixels from the image (when compared to O-Net), although they have *not* been specifically trained for this purpose (i.e. to conduct image denoising). This thus demonstrates a greater propensity of Θ-Net in image denoising applications, though potentially at the expense of incurring network hallucination artifacts.

Retrospectively, further evaluation of the image quality metrics used (namely PSNR, SNR, IMSE & SSIM) seemingly suggests otherwise, with Θ-Net models generally surpassing O-Net model performance when super-resolving DIC micrographs, although the converse occurs for the PCM images. Nonetheless, this assessment proves contrary to the visual discrimination of the features present in the Θ-Net-generated PCM micrographs (when compared with their O-Net counterparts), casting doubt on the veracity of these metrics for quantifying the SR capabilities of a DNN algorithm developed for this purpose. This concern was also surfaced previously in [28], where a suggestion recommending the future development of a suitable metric for quantifying the “super-resolution” quality of an image was proposed, although this metric would probably be difficult to validate, as it would have to consider the illumination type & relative intensity of the light source (in addition to the specimen mounting procedure) being employed, amongst others. In addition, Nyquist sampling criteria would also have to be satisfied [49], to alleviate potential network hallucination due to under-sampling operations which may result in sample noise being confounded as *pseudo*-noise [17].

It would also be noteworthy to mention that the 3D RCAN [25] models used for comparison against the evaluated O-Net & Θ-Net models in this study were developed based on a RGB image spectrally isolated into a 3-channel grayscale image stack within ImageJ 1.52n (NIH, USA), prior to training and executing the models on these images. This was because the 3D RCAN framework [25] was specifically developed to super-resolve grayscale fluorescent micrographs in a 3D Z-stack, although our training images acquired were 2D 24-bit RGB images. The output image stack was subsequently re-merged in ImageJ into a single RGB image.

## Potential Advantages & Limitations of the Present Study

As described in the Discussion section previously, training of the Θ-Net models in the present study was performed intermittently, which resulted in the spikes observed in the loss function plots for the discriminator network (shown in Figures 5 and 7 above). Here (and as was also demonstrated in [50] and [28]), intermittent model training generally results in more accurate models, visualized as the similarity between the generated (output) images & the ground truth/expected image datasets. This phenomenon may be postulated to be due to the minimization of *trapping* of the loss function within local minima in the global error landscape, an aspect which we also surmised in [28].

Nonetheless, a prominent limitation in the current study (which is common to all DNN architectures) refers to the data source which was used to train the models. In the present context, our proposed Θ-Net models were trained on data acquired using a Leica DM4000M microscope with a RisingCam® CMOS camera. Should the model of the microscope and camera used to acquire the test images be different, the models would need to be re-trained, to compensate for the variations in the PSF experienced in the optical train in this respect. For this reason, readers who intend to use our supplied Θ-Net models for their own PCM and/or DIC photomicrographs would have to ensure that an identical microscope and camera setup was used for image acquisition (as highlighted in the present study), or a retraining of the models would be inevitable.

On a separate note, we have also *only* demonstrated the use of Θ-Net in super-resolving PCM and DIC images – two highly popular phase-modulated optical microscopical approaches. Here, and as was also highlighted in [28], numerous other variants of phase microscopy techniques exist, notably Hoffman modulation contrast [51] & oblique illumination [52], as well as the more recently-developed (yet increasingly popular) phase imaging modality – digital holographic microscopy (DHM) [53], amongst others. As we have not verified the applicability of Θ-Net in super-resolving images gleaned from each of these techniques, we are unable to hypothesize on the putative performance of Θ-Net in this respect. It would also be noteworthy to highlight here that these techniques (i.e. PCM & DIC microscopy) are *semi-quantitative* approaches for representing phase variations in the sample, and cannot be used to precisely *quantify* the phasic information due to the non-linear (& thus non-invertible) relationship between the phase and the amplitude in different regions of the sample [54]. However, in our current study, we do not seek to specifically quantify these phase differences, but simply use the said models to translate these phase variations in the sample (caused by the different PSFs in the optical train) into amplitude (brightness) differences for image SR.

Nonetheless, from the results gleaned and presented thus far, we may assuredly infer that the proposed Θ-Net architecture poses significant usability in super-resolving images acquired via PCM and DIC microscopies (two widely utilized label-free diascopic imaging techniques in the optical microscopical space today). In this respect, models developed using Θ-Net hold great promise in future label-free optical nanoscopical applications, facilitating numerous potential extrapolations of engineering applications utilized in the industry today and moving towards the future (i.e. Industry 4.0 & beyond). These may include (but are not restricted to) *bioengineering* applications (for the synthesis of nano-scaffolds to direct proteins to specific sites for proper folding & functional deployment), *materials sciences and electronics engineering* (for quality analysis & cost-friendly nano-scale defect inspection of semiconductor wafers and in nanolithography), as well as in *optical engineering & photonics* applications (to facilitate the research & development of new optical modulators, lasers and diodes).

## Conclusion

In the current study, the performance of our newly-developed DNN architecture (Θ-Net) was checked against other popular state-of-the-art DNN architectures (O-Net [28], 3D RCAN [25], ANNA-PALM [22] & BSRGAN [38]) with respect to its accuracy in super-resolving PCM & DIC micrographs computationally. Our models depict a relatively high level of accuracy, generating images which come close to the ground truth/expected images. Notably, SR images generated by the Θ-Net models of the highly perforated striae of *P. dactylus* var. *dariana* closely resemble the reference SEM micrograph, exemplifying the presence of the poroids in this context. In this regard, one may be led to strongly recommend Θ-Net-derived models for super-resolving PCM and DIC images, although the other compared models (such as 3D RCAN [25]) might perform relatively well for other imaging modalities (such as epi-fluorescence microscopy stacks).

Despite this fact, there exists *no* viable SR metric to be used for objectively quantifying the performance of a DNN model in super resolving images. This dilemma is further exacerbated through the inappropriate use of popular image metrics (such as PSNR, SSIM & IMSE, amongst others) for this purpose in past studies exploring computational SR imaging. A probable resolution of this issue may lie in the development of such a metric *specifically* for image SR, although this would have to consider multiple factors such as the effect of different illumination intensities, camera exposure and gain on the acquired image. A further treatment of this issue is presented in [28] for the interested reader.

Nonetheless, based on our current findings, we can somewhat confidently propose the potentiality of Θ-Net as a means for achieving *in silico* SR for computational phase-modulated nanoscopical applications in the near future, as expounded in the Potential Advantages & Limitations section previously.

## Acknowledgments

The authors would like to extend their sincere appreciations towards past studies conducted by other researchers in the field who have laid some of the groundwork for attaining computational nanoscopy.

## Footnotes

1 Deep-STORM utilizes epi-fluorescence micrographs derived from point source (single molecule) emitters as input images. Here, the dense localization of these fluorescent molecules creates a low-resolution image (akin to that acquired from conventional widefield epi-fluorescence imaging) due to the overlapping emitters (which Deep-STORM seeks to super-resolve in this context).

