## Supplementary Information for "Θ-Net: Achieving Enhanced Phase-Modulated Optical Nanoscopy *in silico* through a computational *‘string of beads’* architecture"

### 1. Technical parameters defined for O-Net & Θ-Net

The architectures for both the O-Net & Θ-Net GANs, as well as the various parameters used for different aspects of these frameworks to super-resolve DIC & phase contrast microscopy (PCM) images in the present study, are illustrated in the following sub-sections:

#### i. O-Net Framework for DIC super-resolution (SR)

The O-Net Pix2Pix GAN to perform DIC SR is depicted in the following diagram:

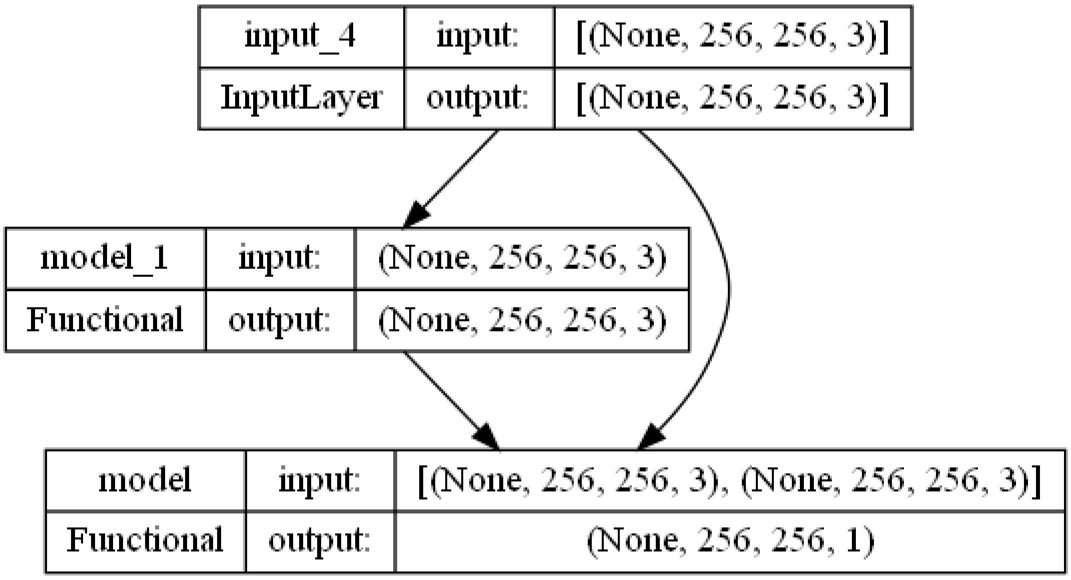


⬤ **Supplementary Figure SF1**: Diagram illustrating the model parameters for the O-Net GAN.

A schematic of the discriminator & generator architectures are provided as follows:

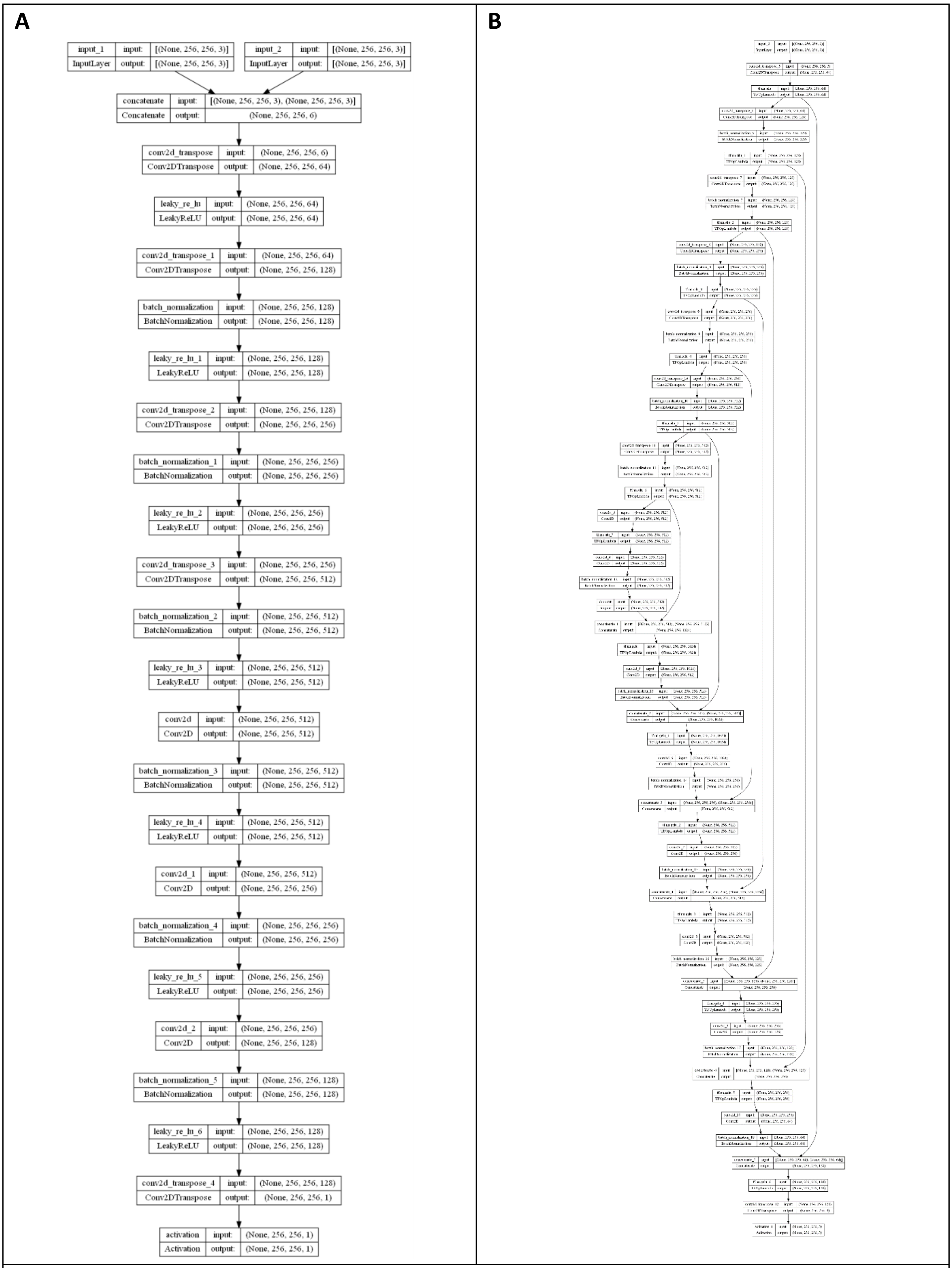


**Supplementary Figure SF2**: Diagram illustrating the model parameters for the O-Net GAN discriminator (**A**) & generator (**B**) respectively.

### ii. Θ-Net Framework for DIC SR

The Θ-Net Pix2Pix GAN used to perform DIC SR in the present study is based on a triple-node string – each node comprising of an O-Net architecture. For the current context, the 1st node of the Θ-Net framework used for DIC SR is identical to that of the O-Net models described in sub-section (i) previously. The 2nd & 3rd nodes of the Θ-Net structure may be described as follows:

<u>For the 2nd Node:</u>

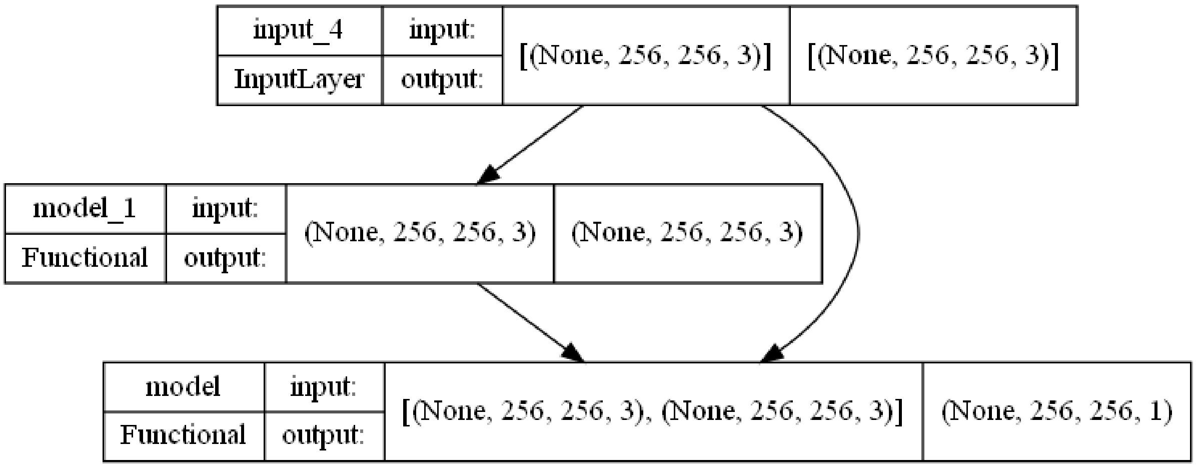


**Supplementary Figure SF3**: Diagram illustrating the model parameters for the 2nd node of the Θ-Net GAN.

As previously, the discriminator & generator architectures may be depicted as follows:

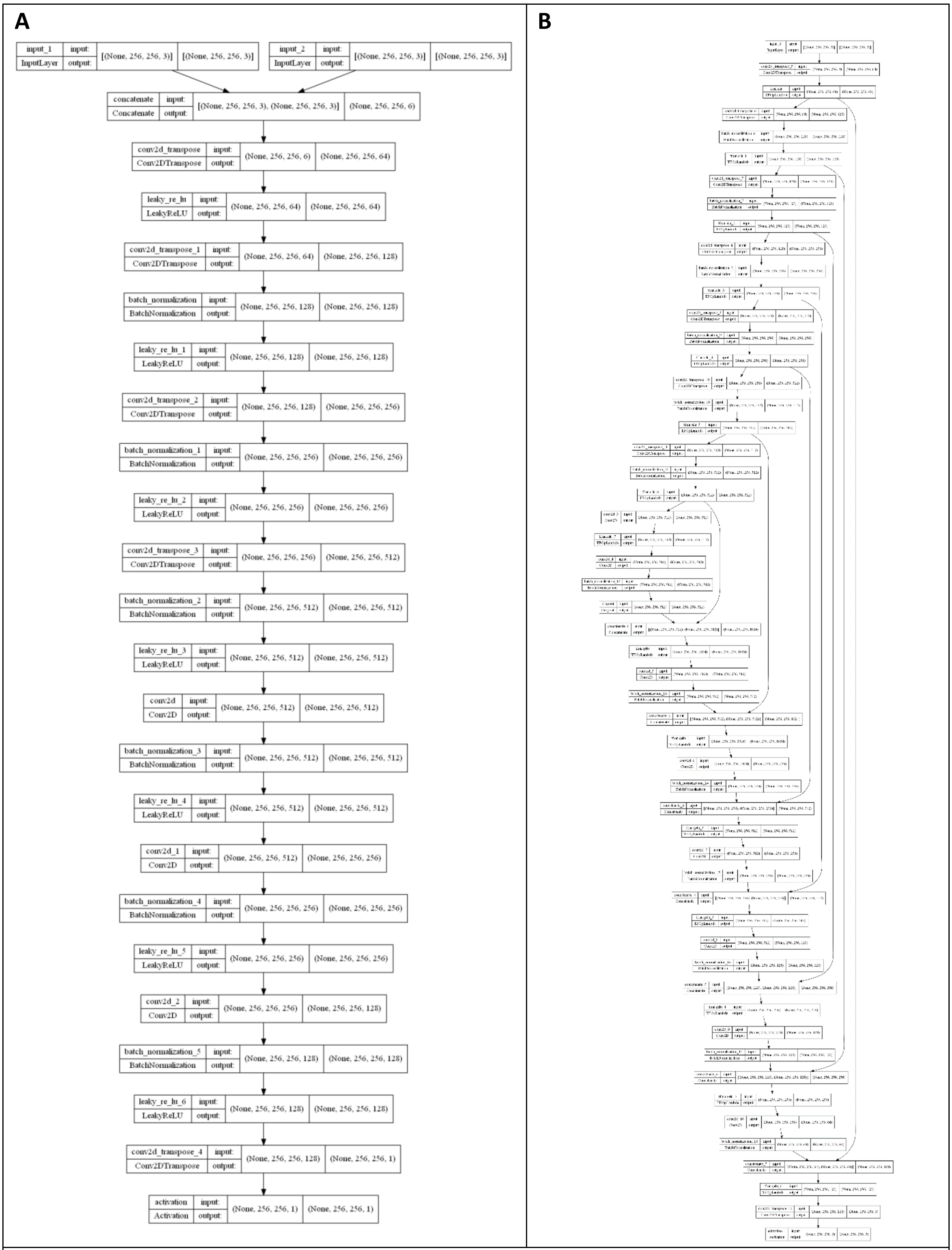


ʘ **Supplementary Figure SF4**: Diagram illustrating the model parameters for the 2nd node Θ-Net GAN discriminator (**A**) & generator (**B**) respectively.

For the 3rd Node:

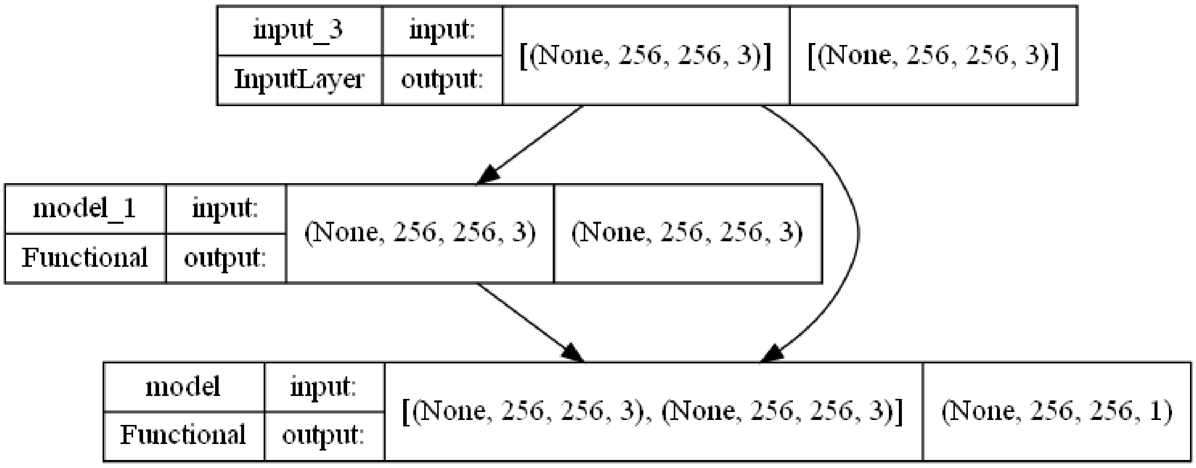


**Supplementary Figure SF5**: Diagram illustrating the model parameters for the 3rd node of the Θ-Net GAN.

In the 3rd node, the discriminator & generator architectures may be depicted as follows:

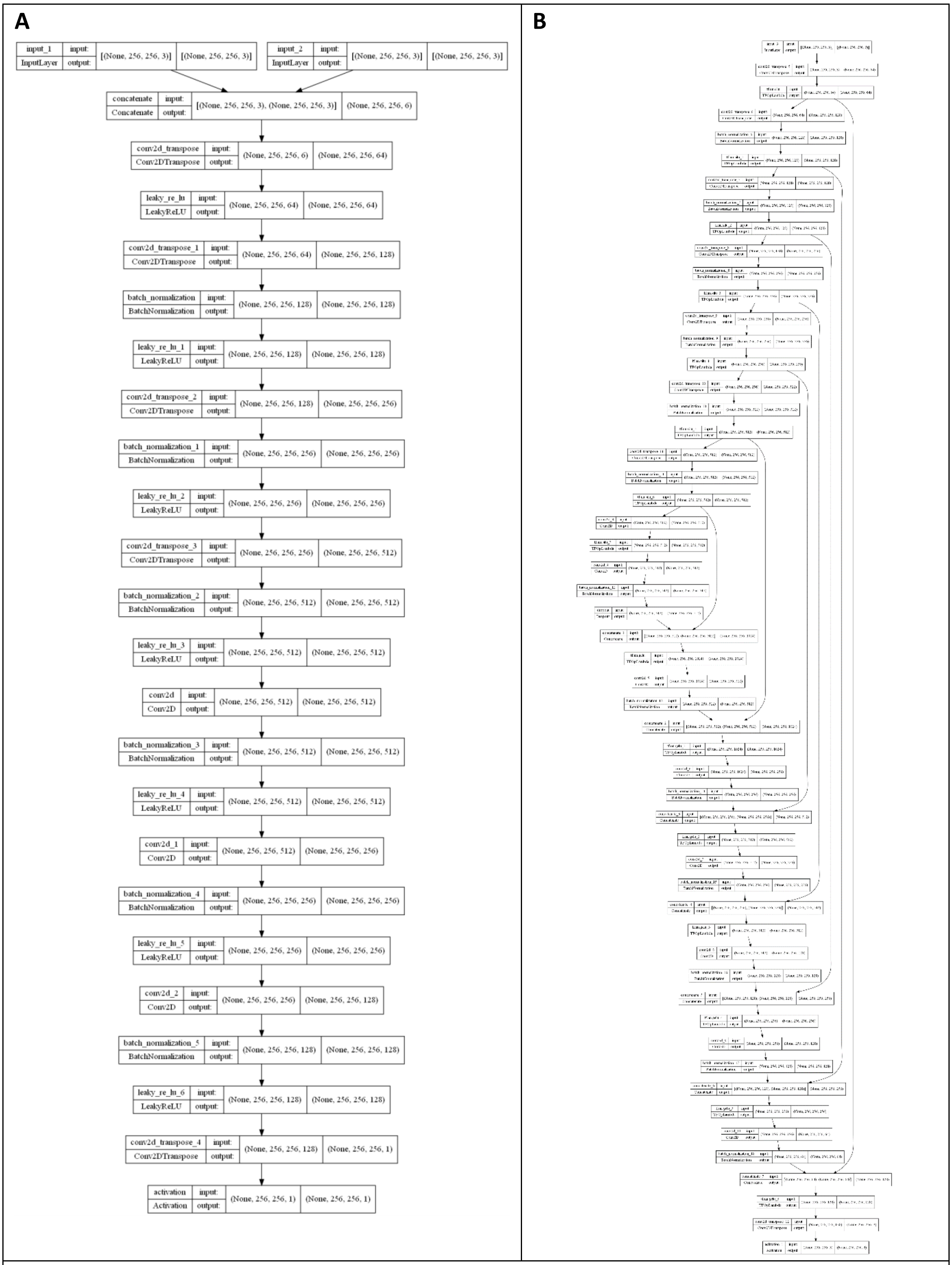


ʘ **Supplementary Figure SF6**: Diagram illustrating the model parameters for the 3rd node Θ-Net GAN discriminator (**A**) & generator (**B**) respectively.

### iii. O-Net Framework for PCM SR

The O-Net Pix2Pix GAN to perform PCM SR is depicted in the following diagram:

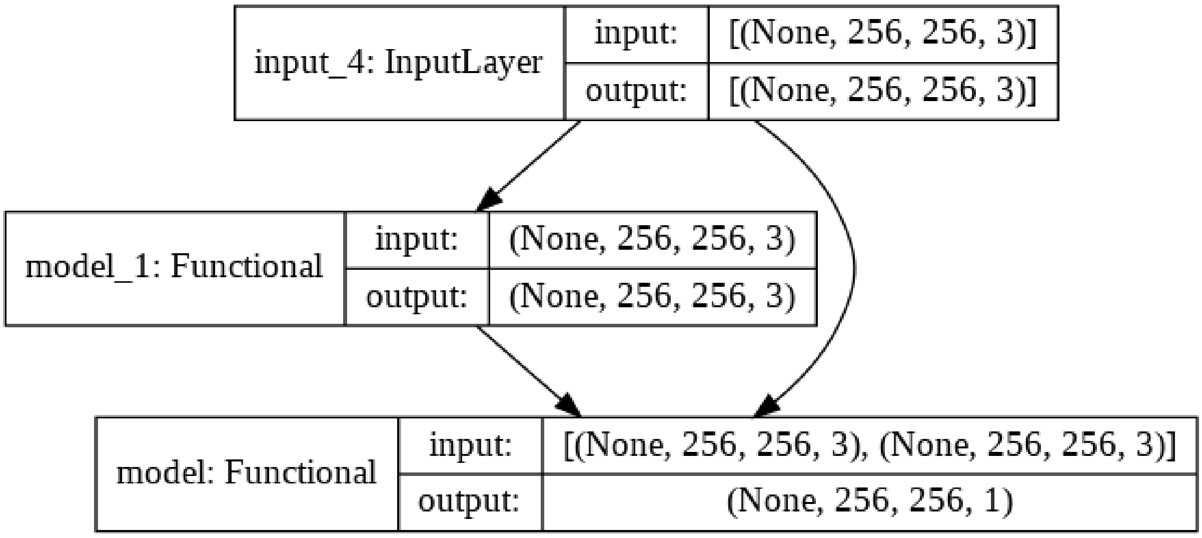


**Supplementary Figure SF7**: Diagram illustrating the model parameters for the O-Net GAN.

A schematic of the discriminator & generator architectures are provided as follows:

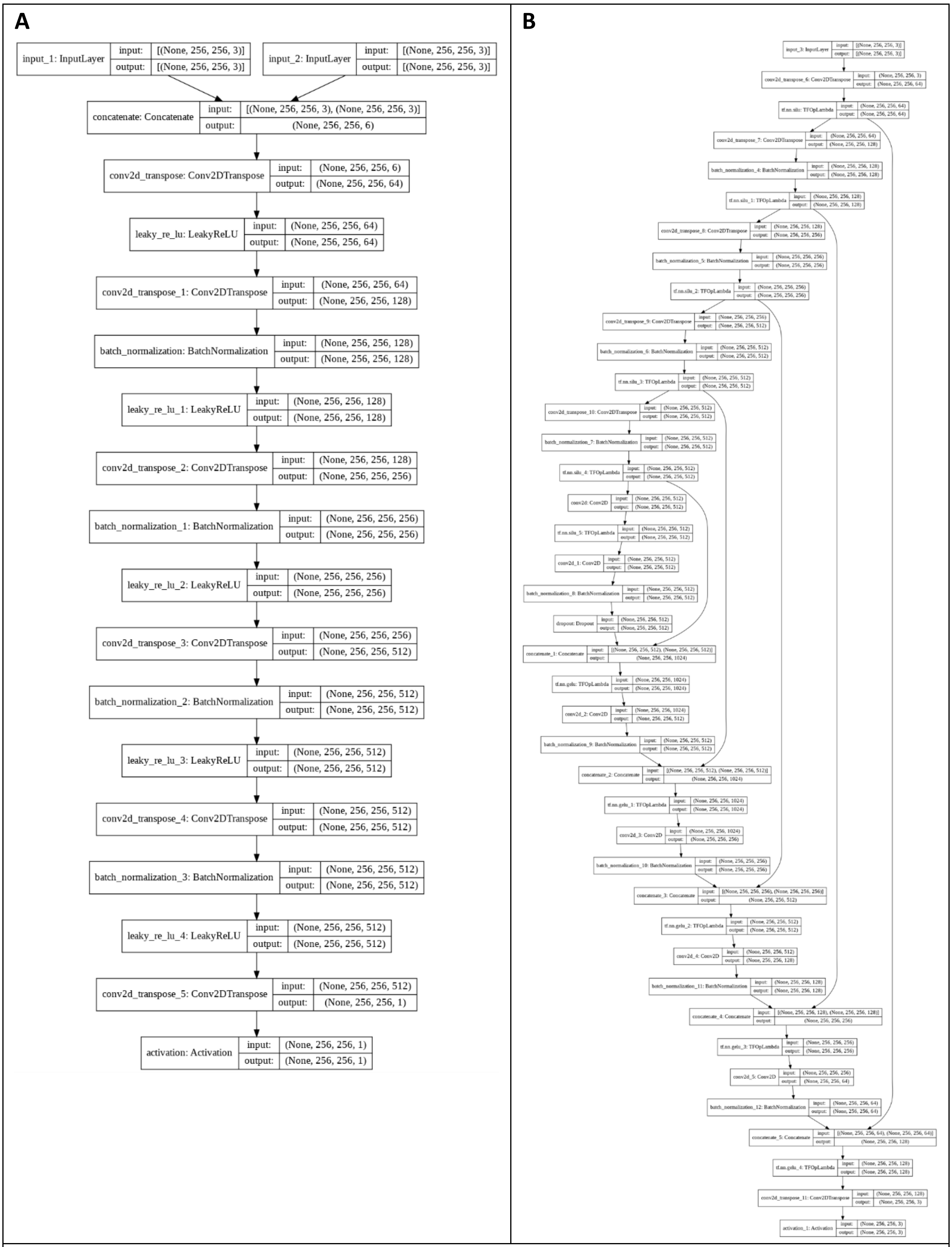


**Supplementary Figure SF8**: Diagram illustrating the model parameters for the O-Net GAN discriminator (**A**) & generator (**B**) respectively.

### iv. Θ-Net Framework for PCM SR

As with the Θ-Net architecture used for DIC SR in sub-section (ii) previously, the Θ-Net Pix2Pix GAN utilized for PCM SR in the current study is also based on a triple-node string, with each node consisting of an O-Net model. However, in the present context, the 1st node of the Θ-Net framework used for PCM SR is identical to that of the O-Net model described in sub-section (iii) previously. In addition, the 3rd node of the Θ-Net structure also employs an identical structure to that of the 3rd node of the Θ-Net framework used for DIC SR [highlighted in subsection (ii) previously]. Thus, only the 2nd node of the Θ-Net string used for PCM SR is unique to this pipeline, with the structure of the said models for this node being described as follows:

<u>For the 2nd Node:</u>

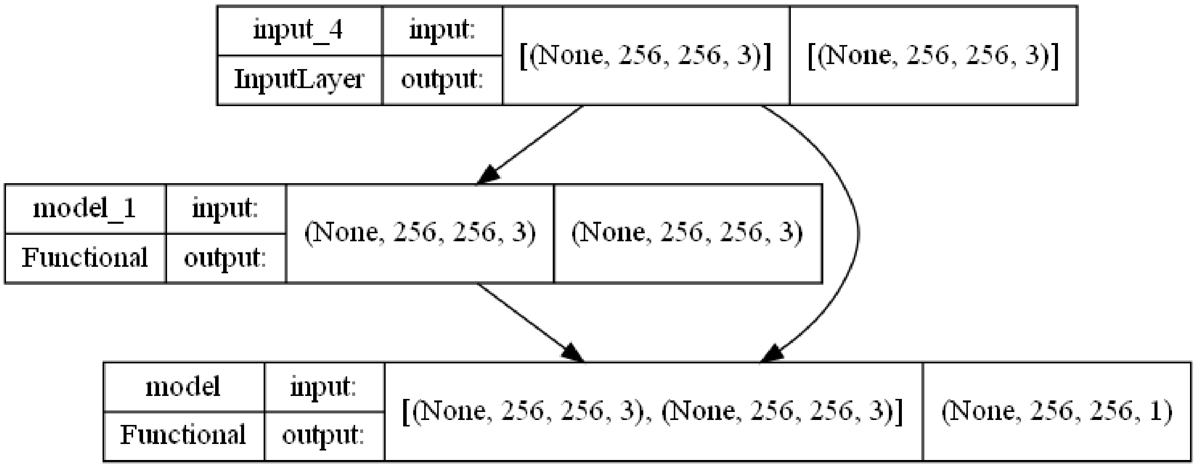


ʘ **Supplementary Figure SF9**: Diagram illustrating the model parameters for the 2nd node of the Θ-Net GAN.

As previously, the discriminator & generator architectures may be depicted as follows:

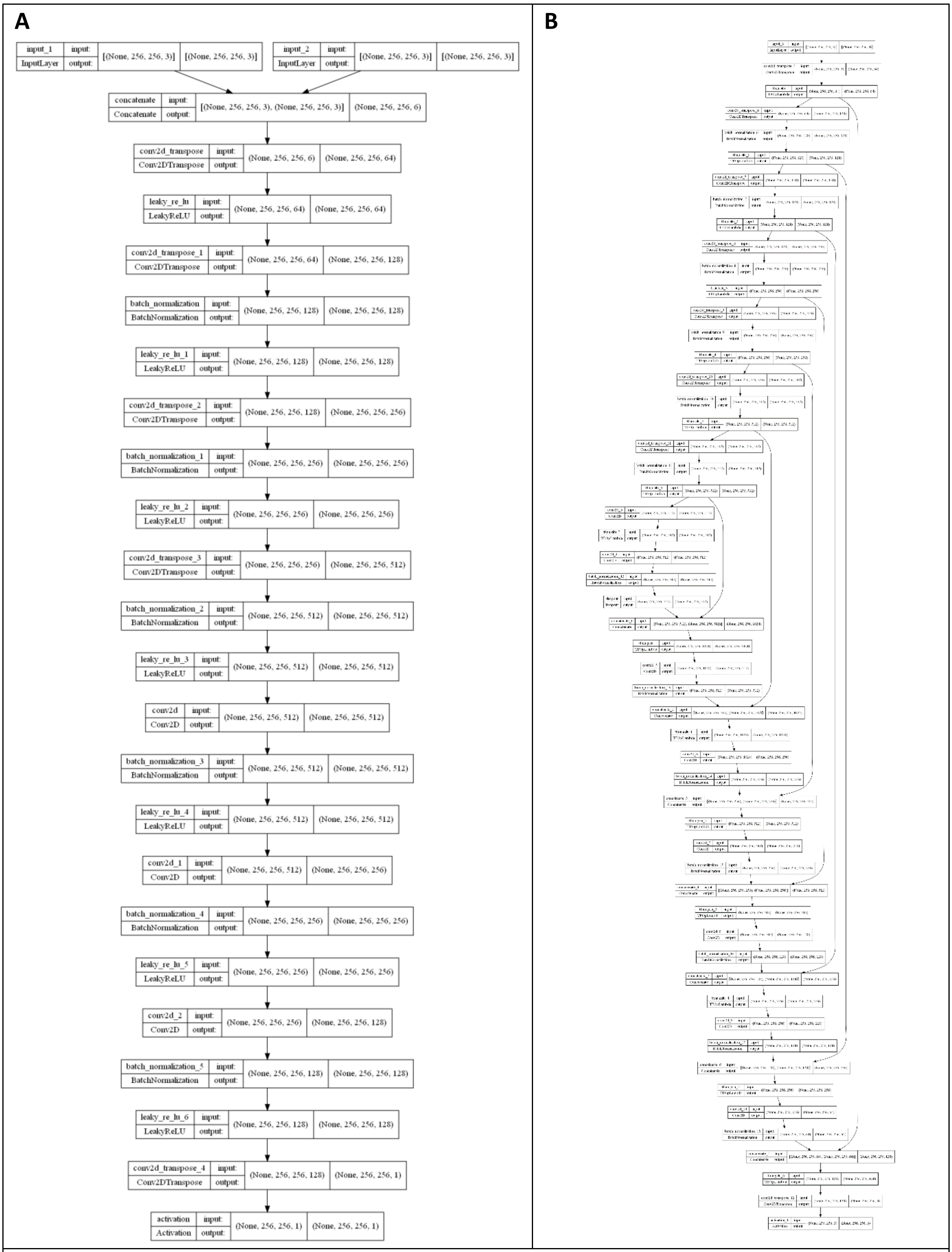


**Supplementary Figure SF10**: Diagram illustrating the model parameters for the 2nd node Θ-Net GAN discriminator (**A**) & generator (**B**) respectively.

### 2. Equations considered for the activation functions

5 different activation functions were used in the O-Net & Θ-Net networks for the present study. These activation functions are as follows:

| Activation Function | Equation | Source |
|---|---|---|
| **Leaky ReLU** | $y = \begin{cases} \alpha \cdot x & x < 0 \\ x & x \geq 0 \end{cases}$ ---------- (SE1) | [1] |
| **Sigmoid** | $y = \frac{1}{1+e^{-x}}$ ------------------------------ (SE2) | [2] |
| **Tanh (Hyperbolic tangent)** | $y = \tanh(x) = \frac{e^x - e^{-x}}{e^x + e^{-x}}$ ------------ (SE3) | [2] |
| **Swish** | $y = \frac{x}{1+e^{-x}}$ ------------------------------ (SE4) | [3] |
| **GELU (Gaussian ELU)** | $y = x\Phi(x)$, where $\Phi(x)$ is the Gaussian CDF of $x$ -------------------- (SE5) | [4] |

### 3. Formulae for the employed image quality metrics

Four image quality metrics are utilized in the current study – the (i) peak signal-noise ratio (PSNR), (ii) signal noise ratio (SNR), (iii) image mean square error (IMSE) & (iv) structural similarity index (SSIM). The formulae underlying each of these metrics are described below:

#### a. Peak signal-noise ratio (PSNR)

PSNR may be mathematically defined as follows:

$$\text{PSNR} = 10\ lg(\frac{\text{PV}^2}{\text{MSE}}) \text{ ---------- (SE6)} \quad \text{(Source: [5])}$$

where PV is the peak (max) value of a pixel in an image (e.g. 255 for an 8-bit image), MSE is the mean squared error and $lg(x) = \log_{10}(x)$ (for some constant $x$). Generally, images having a PSNR >20 imply good (noise-free) images, while those with PSNR values between 18-20 indicate an acceptable image quality standard, although the background of the image is likely polluted with noise.

### b. Signal-noise ratio (SNR)

Another widely-employed image quality metric (akin to PSNR) is SNR, which may be expressed mathematically as follows:

$$\text{SNR} = 10\ lg(\frac{S}{N}) \text{ ------------------- (SE7)}$$

where S and N are the respective strengths of the signal and noise [6]. Here, the MATLAB function **psnr(*A*, *ref*)** also returns the SNR value of the noisy image *A* (with respect to *ref*) [5].

### c. Image mean square error (IMSE)

IMSE represents yet another quantitative approach for comparing 2 images. IMSE may be computed through the following equation (SE8):

$$\text{IMSE}(A, ref) = \sum_{i=1}^{n}(A - ref)^2/n \text{ ----------------- (SE8)}$$

where an image *A* is being compared against a separate reference (*ref*) image of a similar size (*n* is the total number of pixels in either *A* or *ref*). The MATLAB-implementation of IMSE using the in-built function **immse** [7] is being employed in this context.

### d. Structural similarity index (SSIM)

SSIM (the last of the 4 image quality metrics used in the present study) considers 3 factors of imaging, i.e. (i) luminance $\ell$, (ii) contrast $c$ and (iii) structure $s$ [8]. In this respect, SSIM has been determined to be generally superior to other metrics (such as MSE) for comparing different images experiencing varying levels of distortion [9]. Mathematically, SSIM may be expressed as follows (adapted from [8]):

$$\text{SSIM}(A, ref) = [\ell(A, ref)]^{\alpha} \bullet [c(A, ref)]^{\beta} \bullet [s(A, ref)]^{\gamma} \text{ --------------------- (SE9)}$$

where $\mu_A$, $\mu_{ref}$, $\sigma_A$, $\sigma_{ref}$, and $\sigma_{Aref}$ denote the local means, standard deviations, and cross-covariance for images *A, ref* respectively, and

$$\ell(A, ref) = \frac{2\mu_A\mu_{ref} + C_1}{\mu_A^2 + \mu_{ref}^2 + C_1}, \quad c(A, ref) = \frac{2\sigma_A\sigma_{ref} + C_2}{\sigma_A^2 + \sigma_{ref}^2 + C_2}, \quad s(A, ref) = \frac{\sigma_{Aref} + C_3}{\sigma_A\sigma_{ref} + C_3} \text{ ------------ (SE10)}$$

(Source: [8])

By default, MATLAB assigns $\alpha = \beta = \gamma = 1$ and $C_3 = C_2 / 2$, thereby reducing SSIM to the following:

$$\text{SSIM}(A, ref) = \frac{(2\mu_A\mu_{ref} + C_1)(2\sigma_{Aref} + C_2)}{(\mu_A^2 + \mu_{ref}^2 + C_1)(\sigma_A^2 + \sigma_{ref}^2 + C_2)} \text{ ---------------------- (SE11)}$$ (Source: [8])

Computation of these four metrics for the images analyzed in the present study were performed in MATLAB R2022B (© 1984-2022, The MathWorks, Inc).

### 4. Models, Codes & Figures Availability

The models, codes and images utilized in the present study may be downloaded via the link below:

**Download link:**

https://bit.ly/3Osr4WC

All codes are written in Python or MATLAB – the former having the **.py** extension, while the latter has the **.m** extension. It is recommended that the reader intending to execute the supplied codes for validation use Python⩾3.8 and MATLAB ⩾ R2020a to do so. The model files used for the generating the images depicted in the present study have a **.h5** extension.
